# Proteomic studies of the Arabidopsis TRAPP complexes reveal conserved organization and a novel plant-specific component with a role in plant development

**DOI:** 10.1101/684258

**Authors:** Veder J. Garcia, Shou-Ling Xu, Raksha Ravikumar, Wenfei Wang, Liam Elliott, Mary Fesenko, Melina Altmann, Pascal Falter-Braun, Ian Moore, Farhah F. Assaad, Zhi-Yong Wang

**Affiliations:** Department of Plant Biology, Carnegie Institution for Science, Stanford, CA, 94305; Plant Science Department, Botany, Technische Universität München, 85354 Freising, Germany; Basic Forestry and Proteomics Research Center, Haixia Institute of Science and Technology, Fujian Agriculture and Forestry University, Fuzhou, China; Department of Plant Sciences, South Parks Road, Oxford, OX1 3RB, UK; Institute of Network Biology (INET), Helmholtz Zentrum München, Deutsches Forschungszentrum für Gesundheit und Umwelt (GmbH), 85764 Neuherberg, Germany; Faculty of Biology, Microbe-Host-Interactions, Ludwig-Maximilians-Universität (LMU) München, 82152 Planegg-Martinsried, Germany

## Abstract

How the membrane trafficking system spatially organizes intracellular activities and intercellular signaling networks is not well understood in plants. The Transport Protein Particle (TRAPP) complexes are known to play key roles in selective delivery of membrane vesicles to various subcellular compartments in yeast and animals, but remain to be fully characterized in plants. Here we interrogate the TRAPP complexes in Arabidopsis using quantitative proteomic approaches. TRS33 is a component shared by all TRAPP complexes in yeast and animals, and the Arabidopsis AtTRS33 is essential for the subcellular dynamics of other TRAPP components. Affinity purification of AtTRS33 followed by quantitative mass spectrometry identified fourteen interacting proteins; these include not only thirteen homologs of all known TRAPP components in yeast and mammals but also a novel protein we named TRAPP-interacting plant protein (TRIPP), which is conserved in multi-cellular photosynthetic organisms. Proteomic and molecular analyses showed that TRIPP specifically associates with the TRAPPII complex *in vivo* and directly interacts with the TRAPPII-specific subunits but not the subunits shared with TRAPPIII. TRIPP co-localizes with a subset of TRS33 compartments, and its localization is disrupted in the *trs33* mutant. Loss-of-function *tripp* mutation caused growth and reproductive development defects, including partial photomorphogenesis in the dark. Our study demonstrates that plants possess at least two distinct TRAPP complexes similar to metazoan, and identifies TRIPP as a novel plant-specific component of the TRAPPII complex with important functions in plant growth and development.

## Introduction

Vesicular protein trafficking is a highly organized and regulated cellular process that ensures delivery of proteins to their correct subcellular compartments (Rosquete and Drakakaki, 2018). Vesicle transport spatially organizes not only intracellular structures and metabolic activities, but also intercellular signaling systems that control development. Vesicle transport involves numerous protein complexes and is regulated at multiple stages, starting with vesicle budding from a donor membrane and ending with the fusion stage, where a vesicle merges with a specific acceptor membrane (Brocker et al., 2010). Prior to the fusion stage, tethering factors initiate and maintain specific contact between donor and acceptor membranes to hold the vesicle in close proximity to the target membrane (Whyte and Munro, 2002). Thus, tethering factors play key role in organizing vesicle trafficking. However, the functions of tethering factors in plant development remain largely uncharacterized.

In eukaryotes, there are two broad classes of tethering factors, long coiled-coil proteins and multi-subunit tethering complexes (MTCs) (Brocker et al., 2010; Yu and Hughson, 2010; Ravikumar et al., 2017; Takemoto et al., 2018). Coiled-coil tethers are long, dimeric proteins that are found mainly on the Golgi and early endosomes (Lurick et al., 2018), while MTCs contain several subunits in a modular form and are located on organelles throughout the secretory and endocytic pathways. A well-studied MTC in yeast and animals is the Transport Protein Particle (TRAPP) complex, which is involved in Endoplasmic Reticulum (ER) to Golgi and Golgi-mediated trafficking as well as in autophagy (Barrowman et al., 2010; Scrivens et al., 2011; Vukasinovic and Zarsky, 2016). In yeast, mutations in TRAPPs are typically lethal or cause strongly impaired growth (Kim et al., 2016). In humans, mutations in TRAPP components have been implicated in a variety of diseases (Milev et al., 2015; Sacher et al., 2018).

TRAPP complexes can exist in a variety of modular forms (Robinett et al., 2009; Kim et al., 2016; Riedel et al., 2018). In yeast and metazoan, two TRAPP complexes (II and III) composed of shared and complex-specific subunits have been confirmed (Barrowman et al., 2010; Lynch-Day et al., 2010; Thomas et al., 2018). The metazoan TRAPP complexes contain orthologues of all yeast TRAPP subunits, but have also evolved additional subunits and rearrangement of complex composition. The TRAPPIII complex contains two additional metazoan-specific subunits, TRAPPC11 and TRAPPC12 (Scrivens et al., 2011; Zhao et al., 2017). Tca17 and Trs65 are TRAPPII-specific subunits in yeast, but their metazoan orthologues (TRAPPC2L and TRAPPC13, respectively) are found in TRAPPIII complexes. The subunit composition of TRAPP complexes specifies localization and function. In yeast, TRAPPII functions in late Golgi and TRAPPIII in early Golgi and autophagy (Kim et al., 2016; Riedel et al., 2018; Thomas et al., 2018). In metazoans, TRAPPII is involved in membrane traffic in the biosynthetic pathway, and TRAPPIII functions in autophagosome formation and autophagy (Yamasaki et al., 2009; Scrivens et al., 2011; Bassik et al., 2013; Lamb et al., 2016; Ramirez-Peinado et al., 2017; Sacher et al., 2018). Furthermore, TRAPP complexes have been shown to function as Guanine nucleotide Exchange Factors (GEF) for Rab GTPases in both yeast and metazoans (Jenkins et al., 2018; Riedel et al., 2018; Thomas et al., 2018; Thomas et al., 2019).

In plants, TRAPP complex formation and modularity remain to be elucidated. Although the Arabidopsis genome encodes orthologous TRAPP components, evidence as to the existence of most TRAPP components at the protein level and whether or how these assemble into TRAPP complexes is incomplete. Two Arabidopsis TRAPP subunits have been characterized to some extent in plants; AtTRS120 and AtTRS130 (Jaber et al., 2010; Thellmann et al., 2010; Naramoto et al., 2014; Rybak et al., 2014). Both AtTRS120/VAN4 and AtTRS130/CLUB are required for cell plate biogenesis. Mutations in *AtTRS120* or *AtTRS130* cause seedling lethality and canonical cytokinesis⍰ defective phenotypes, including cell wall stubs and incomplete cross walls (Jaber et al., 2010; Thelmann et al., 2010; Qi et al., 2011). Also, in both mutants, vesicles accumulate at the equator of dividing cells but fail to assemble into cell plates (Jaber et al., 2010; Thellmann et al., 2010; Rybak et al., 2014; Ravikumar et al., 2017). In accordance with TRAPP localization in other organisms, AtTRS120 and AtTRS130 localize at the *trans*-Golgi Network (TGN) (Qi et al., 2011; Naramoto et al., 2014; Ravikumar et al., 2017; Ravikumar et al., 2018). Recently, in a genetic characterization, the Arabidopsis orthologue of the yeast/metazoan shared subunit BET5 has been shown to be essential for pollen exine pattern formation in Arabidopsis (Zhang et al., 2018).

The above-mentioned studies provide strong evidence for critical functions of TRAPP components in plants. Previous sequence analysis and proteomic studies have predicted and identified partial TRAPP components, respectively (Drakakaki et al., 2012; Paul et al., 2014; Rybak et al., 2014). However, experimental evidence is lacking with regard to whether plants form the same number of TRAPP complexes as yeast and metazoans, what the subunit composition of such complexes might be and whether there are plant-specific TRAPP components. To address these questions, we have taken a holistic proteomic approach to document the subunit composition and interactors of TRAPP complexes in Arabidopsis. We identified eleven subunits that are conserved in yeast and metazoans, and two that are only conserved between metazoans and plants. Additionally, we identify and characterize a novel TRAPP interacting plant protein (TRIPP) that is present only in multicellular photosynthetic organisms. Genetic evidence shows that TRIPP plays an important role in plant growth, reproduction, and light-dependent development. Our study demonstrates that Arabidopsis TRAPP components form similar complexes as reported in metazoans. The study also uncovers a novel plant-specific TRAPPII interactor that plays important roles in plant growth and development.

## RESULTS

### AtTRS33 is required for the membrane association of a TRAPPII-specific subunit

Previous studies of TRAPP subunits in plants have partially revealed TRAPP components, for instance, Drakakaki and her colleagues identified five TRAPP subunits from isolated SYP61 TGN compartments in Arabidopsis using mass spectrometric analysis (Drakakaki et al., 2012). A more recent AtTRS120 IP-MS study identified five subunits, four overlapping with the results of Drakakaki et al. (Rybak et al., 2014). However, a computational study that used OrthoMCL and PGAP algorithms to find vesicular transport orthologues in yeast and plant genomes, identified at least nine TRAPP components encoded in Arabidopsis (Paul et al., 2014).

To further identify protein components of TRAPP complexes in plants, we carried out quantitative mass spectrometry analysis of proteins associated with TRS33, a subunit known to be shared by different TRAPP complexes studied in both yeast and metazoans (Kim et al., 2016; Riedel et al., 2018). We reasoned that an analysis of such a shared subunit would give us a more comprehensive view of TRAPP complexes in Arabidopsis than has thus far been available from studies based on TRAPPII-specific subunits alone (Rybak et al., 2014; Ravikumar et al., 2018). In Arabidopsis, AtTRS33 has been implicated in cytokinesis and found in the TRAPPII interactome (Thellmann et al., 2010; Rybak et al., 2014).

Prior to using TRS33 to further study TRAPP composition, we first sought to confirm TRS33’s functional relationship to TRAPPII in Arabidopsis. To this end, we monitored the effect of *trs33*-*1* mutation on the localization dynamics of the TRAPPII-specific subunit AtTRS120. In the wild type, the TRS120-GFP fusion protein localizes to both the cytosol and to TGN compartments in interphase cells (Figure 1A; Rybak et al., 2014; Ravikumar et al., 2018). In contrast, in the *trs33-1* mutant, the signal was only observed in the cytosol, and no endomembrane compartments were detected (Figure 1A). This is reminiscent of the mis-localization of a TRAPPII-specific subunit (Trs130-GFP) to the cytosol in *trs33* mutants of yeast (Tokarev et al., 2009). At early cytokinetic stages, TRS120-GFP clearly localized to the cell plate in the wild type, but was present as a diffuse cytosolic cloud around the cell plate in *trs33-1* (Figure 1B, 1D). Furthermore, at the end of cytokinesis, TRS120-GFP re-localized to the leading edges of the cell plate in the wild type but was, at best, visible as a weak and relatively diffuse signal at the leading edges of *trs33-1* expanding cell plates (Figure 1C). Thus, AtTRS33 is required for the membrane association of AtTRS120 and for its proper localization dynamics during cytokinesis. This establishes a clear functional link between AtTRS33 and the Arabidopsis TRAPPII complex.

**Figure 1.**
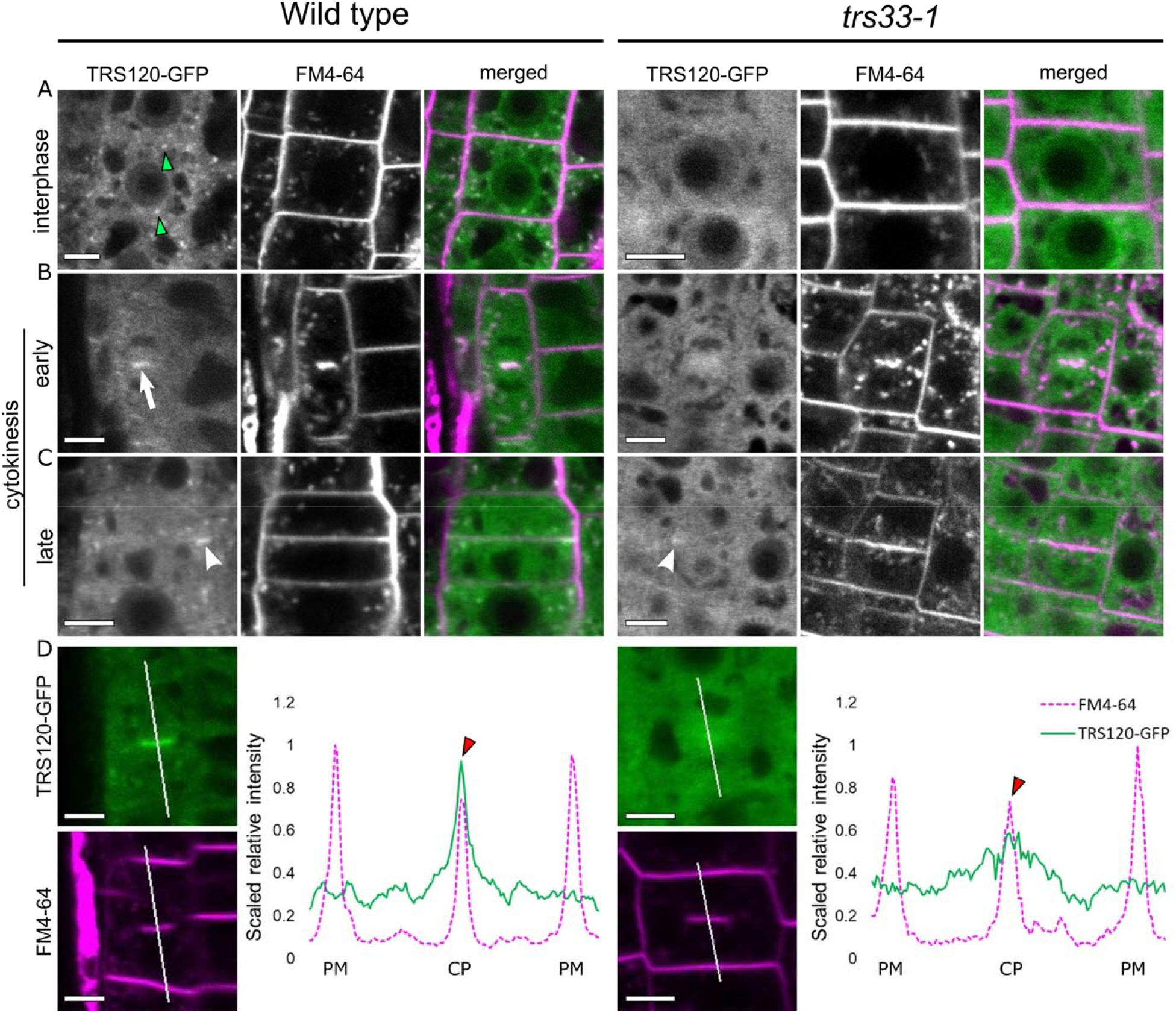
TRS33 is required for normal subcellular localization of TRS120-GFP. Live imaging of TRS120-GFP (green) and FM4-64 (magenta in the merged panels) in roots of *TRS120:TRS120-GFP* plants. CP cell plate. PM: plasma membrane. **A.** Cells at interphase show TRS120-GFP enriched at endomembrane compartments in the wild type (green arrowheads) but not in *trs33-1*, where only a cytosolic haze can be seen. **B.** At early cytokinetic stages (CP initiation and biogenesis), TRS120-GFP is enriched at the cell plate (white arrow) in the wild type but not in *trs33-1* **C.** At late cytokinetic stages (CP insertion and maturation), TRS120-GFP reorganizes to the leading edges of the cell plate (white arrowhead) in the wild type. In contrast, only a weak and relatively diffuse TRS120-GFP signal can be detected at the leading edges of the cell plate in *trs33-1* mutants (white arrowhead). **D.** Line graphs depicting scaled relative fluorescence intensities. A sharp peak is seen at the CP in the wild type (red arrowhead) but not in *trs33-1*. At least ten seedlings were imaged per marker line. n = 8 wild type, n = 7 *trs33-1* cytokinetic cells. Size bars = 5μm.

### Quantitative mass spectrometry characterization of the TRAPP complexes

To identify proteins associated with AtTRS33, we used <u>s</u>table <u>i</u>sotope <u>l</u>abeling <u>i</u>mmuno<u>p</u>recipitation followed by quantitative <u>m</u>ass <u>s</u>pectrometry (SILIP-MS). Transgenic Arabidopsis seedlings expressing AtTRS33 fused with Myc and His tags from the native *AtTRS33* promoter (*TRS33:TRS33-MycHis*) in *trs33-1* mutant background were grown for 14 days on medium containing light (^14^N) nitrogen, along with wild type as control grown on heavy (^15^N) nitrogen. The isotopes were switched in the replicate experiment. Root and shoot tissues were harvested separately. Each pair of ^14^N- and ^15^N-labelled sample and control were mixed together before immunoprecipitation with anti-Myc antibody beads (Figure 2A). Immunoprecipitated proteins were separated by SDS-PAGE, in-gel digested and analyzed in an Orbitrap mass spectrometer. Enrichment by TRS33 was quantified by the ^14^N/^15^N ratios of the identified peptides. MS analysis identified over 1000 proteins in each experiment, but only proteins that showed >2-fold enrichment in the TRS33-MycHis samples over the controls in both forward and reciprocal labeling replicates were considered as TRS33 interactors. We identified fourteen TRS33-interacting proteins in samples from both roots and shoots (Table I). SILIP-MS using transgenic Arabidopsis plants that overexpress TRS33-YFP also identified the same 14 interacting proteins (Supplemental Table I). These include all orthologues of the shared TRAPP subunits reported in yeast and metazoans (TRS23, TRS33, TRS31, BET3, BET5, and TRS20) and of the complex-specific subunits (TCA17, TRS120, TRS130, TRS85, TRAPPC11, TRAPC12, and TRAPPC13) (Riedel et al., 2018; Sacher et al., 2018). Intriguingly, the identified TRS33-associated proteins include an additional protein that has no orthologues in yeast or metazoans (Figure 2B, Table I, Supplemental Table I, and Supplemental Figure 1). This protein is composed of 838 amino acids and is listed as unknown in The Arabidopsis Information Resource (TAIR) database. We named this protein <u>TR</u>APP <u>I</u>nteracting <u>P</u>lant <u>P</u>rotein (TRIPP). Taken together, the mass spectrometry results demonstrate that plant TRAPP complexes contain not only homologs of all known TRAPP components in yeast and metazoans, but also a plant-specific component TRIPP.

**Figure 2.**
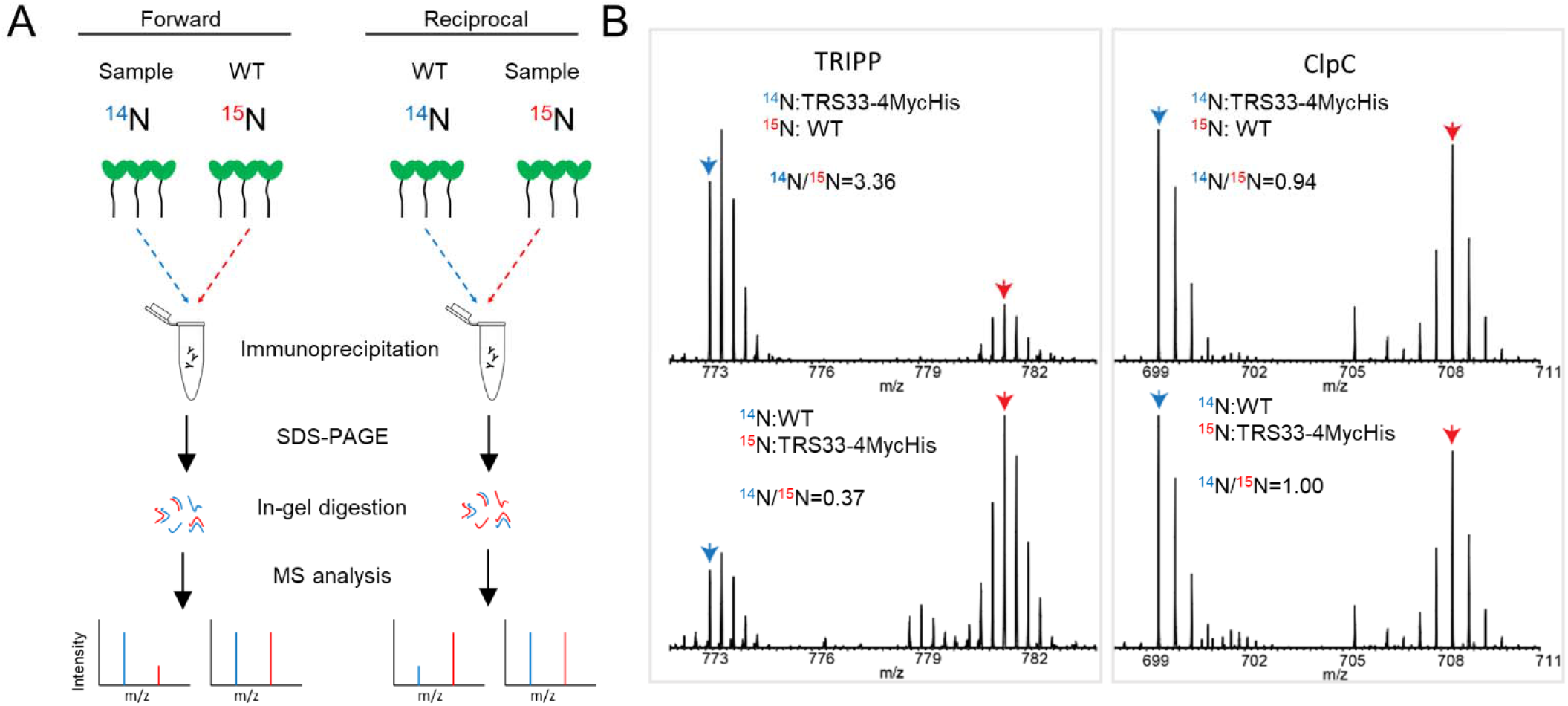
TRAPP-associated proteins were identified by Immunoprecipitation of TRS33. **A.** Experimental work flow for the <u>S</u>table <u>I</u>sotope <u>L</u>abeling <u>I</u>mmunoprecipitation <u>M</u>ass <u>S</u>pectrometry (SILIP-MS) in Arabidopsis. **B.** MS1 spectra show the enrichment of TRIPP in SILIP-MS of TRS33:TRS33-4MycHis/*trs33-1* versu Wild type (Col-0) control, and no enrichment of ClpC, a non-specific protein. Blue and red rows point to monoisotopic peaks of ^14^N- and ^15^N-labeled peptides, respectively.

**Table I.** Fourteen Arabidopsis TRAPP components were identified by stable isotope labeling immunoprecipitation mass spectrometry (SILIP-MS). SILIP-MS was performed using the *TRS33*:*TRS33*-*4MycHis*/*trs33-1* complemented plants (Root and Shoot separately). Number of unique peptides (Peptide No.), percentage of sequence coverage (% Cov), and ratio of signal intensities between samples labelled with light (L, ^14^N) and heavy (H, ^15^N) isotopes (L/H ratio) are shown for two biological replicates, in which either the Col-0 control (Forward) or the TRS33-MycHis sample (Reciprocal) was labelled with ^15^N. ^a^Subunits previously identified in a proteomic study using isolated TGN compartments (Drakaki et. al, 2012). ^b^Subunits previously identified in label-free IP-MS of AtTRS130 (Rybak et al 2014). Yeast and human orthologue names are based on published work (Thomas et al., 2018 and Riedel et al, 2018, respectively).

| TRAPP subunits identified by TRS33 Immunoprecipitation from *Arabidopsis* roots |  |  |  |  |  |  |  | Yeast Orthologue | Human Orthologue |
| --- | --- | --- | --- | --- | --- | --- | --- | --- | --- |
|  | Forward |  |  | Reciprocal |  |  |  |  |  |
| Gene ID | Peptide No. | % Cov | L/H ratio | Peptide No. | % Cov | L/H ratio | Protein Name |  |  |
| AT1G51160 | 5 | 41.4 | 4.47 | 4 | 28.4 | 0.41 | BET5 | BET5 | TRAPPC1 |
| AT1G80500 | 3 | 28.9 | 4.13 | 3 | 28.9 | 0.29 | TRS20 | TRS20 | TRAPPC2 |
| AT2G20930 | 6 | 52.9 | 3.79 | 5 | 52.9 | 0.39 | TCA17 | TCA17 | TRAPPC2L |
| AT5G54750 | 12 | 55.4 | 11.4 | 12 | 54.3 | 0.18 | BET3<sup>a,b</sup> | BET3 | TRAPPC3 |
| AT5G02280 | 5 | 42.6 | 2.72 | - | - | - | TRS23 | TRS23 | TRAPPC4 |
| AT5G58030 | 5 | 22.6 | 2.48 | 2 | 10.3 | 0.37 | TRS31<sup>a,b</sup> | TRS31 | TRAPPC5 |
| AT3G05000 | 11 | 61.3 | 7.7 | 11 | 60.7 | 0.07 | TRS33<sup>a,b</sup> | TRS33 | TRAPPC6 |
| AT5G16280 | 16 | 11.7 | 7.27 | 13 | 9.5 | 0.15 | TRS85<sup>a</sup> | TRS85 | TRAPPC8 |
| AT5G11040 | 32 | 32.7 | 3.62 | 23 | 25.6 | 0.40 | TRS120/VAN4<sup>a,b</sup> | TRS120 | TRAPPC9 |
| AT5G54440 | 33 | 31 | 4.19 | 13 | 12.3 | 0.44 | TRS130/CLUB<sup>b</sup> | TRS130 | TRAPP10 |
| AT5G65950 | 20 | 17.6 | 7.76 | 12 | 11.2 | 0.18 | TRAPPC11 | - | TRAPPC11 |
| AT4G39820 | 15 | 35.8 | 8.06 | 8 | 21.3 | 0.24 | TRAPPC12 | - | TRAPPC12 |
| AT2G47960 | 14 | 37.1 | 2.84 | 14 | 33.3 | 0.52 | TRS65 | TRS65 | TRAPPC13 |
| AT3G17900 | 22 | 30.4 | 3.54 | 19 | 24.8 | 0.39 | TRIPP | - | - |
| TRAPP subunits identified by TRS33 Immunoprecipitation from *Arabidopsis* shoots |  |  |  |  |  |  |  | Yeast orthologue | Human Orthologue |
|  | Forward |  |  | Reciprocal |  |  |  |  |  |
| Gene ID | Peptide No. | % Cov | L/H ratio | Peptide No. | % Cov | L/H ratio | Protein Name |  |  |
| AT1G51160 | 4 | 34.9 | 14.7 | 2 | 14.2 | 0.16 | BET5 | BET5 | TRAPPC1 |
| AT1G80500 | 1 | 8.9 | 12.9 | 2 | 20 | 0.10 | TRS20 | TRS20 | TRAPPC2 |
| AT2G20930 | 6 | 63.6 | 3.9 | 4 | 36.4 | 0.36 | TCA17 | TCA17 | TRAPPC2L |
| AT5G54750 | 12 | 55.4 | 30.2 | 11 | 46.8 | 0.07 | BET3<sup>a,b</sup> | BET3 | TRAPPC3 |
| AT5G02280 | 6 | 51.8 | 7.1 | - | - | - | TRS23 | TRS23 | TRAPPC4 |
| AT5G58030 | 4 | 29.2 | 11.9 | 6 | 29.7 | 0.14 | TRS31<sup>a,b</sup> | TRS31 | TRAPPC5 |
| AT3G05000 | 12 | 74.0 | 37.9 | 8 | 52.6 | 0.04 | TRS33<sup>a,b</sup> | TRS33 | TRAPPC6 |
| AT5G16280 | 16 | 12.7 | 10.3 | 13 | 8.6 | 0.12 | TRS85<sup>a</sup> | TRS85 | TRAPPC8 |
| AT5G11040 | 24 | 22.5 | 5.5 | 19 | 18.6 | 0.23 | TRS120/VAN4<sup>a,b</sup> | TRS120 | TRAPPC9 |
| AT5G54440 | 23 | 20.2 | 4.7 | 18 | 17.1 | 0.27 | TRS130/CLUB<sup>b</sup> | TRS130 | TRAPP10 |
| AT5G65950 | 27 | 22.8 | 12.8 | 13 | 12.4 | 0.12 | TRAPPC11 | - | TRAPPC11 |
| AT4G39820 | 9 | 24.3 | 15.3 | 10 | 24.3 | 0.11 | TRAPPC12 | - | TRAPPC12 |
| AT2G47960 | 14 | 32.4 | 5.2 | 12 | 30.5 | 0.26 | TRS65 | TRS65 | TRAPPC13 |
| AT3G17900 | 19 | 28.6 | 3.8 | 17 | 21.1 | 0.32 | TRIPP | - | - |

### TRIPP behaves like a component of the TRAPP II complex in SILIP-MS experiments

To confirm that TRIPP associates with TRAPP components and to determine whether it is specific to a given TRAPP complex, we performed an additional SILIP-MS using TRIPP as bait. Transgenic plants expressing TRIPP-YFP under a 35S promoter, which complemented a *tripp-1* mutant, were grown along non-transgenic wild type as control, as described above for TRS33 SILIP-MS experiment. The TRIPP-YFP and associated proteins were Immunoprecipitated with anti-GFP antibody, and analyzed by mass spectrometry (Figure 2A). The SILIP-MS data showed unambiguously that TRIPP pulled down nine TRAPP subunits (Figure 3A, Supplemental Table II). These include all six TRAPP shared subunits (TRS23, TRS33, TRS31, BET3, BET5, TRS20) and three TRAPPII-specific subunits (TCA17, TRS120 and TRS130) (Figure 3A, 3B, 3C and Supplemental Table II). Four TRAPP subunits (TRS85, TRAPPC11, TRAPPC12 and TRAPPC13) identified in the TRS33 SILIP-MS showed no interaction with TRIPP in these SILIP-MS experiments. None of these proteins were detected in both replicate experiments or enriched by TRIPP-YFP in either single experiment. Interestingly, these are homologs of the subunits specific to TRAPPIII in animals, suggesting that Arabidopsis similarly possesses distinct TRAPPII and TRAPPIII complexes, and that TRIPP specifically associates with the plant TRAPPII complex.

**Figure 3.**
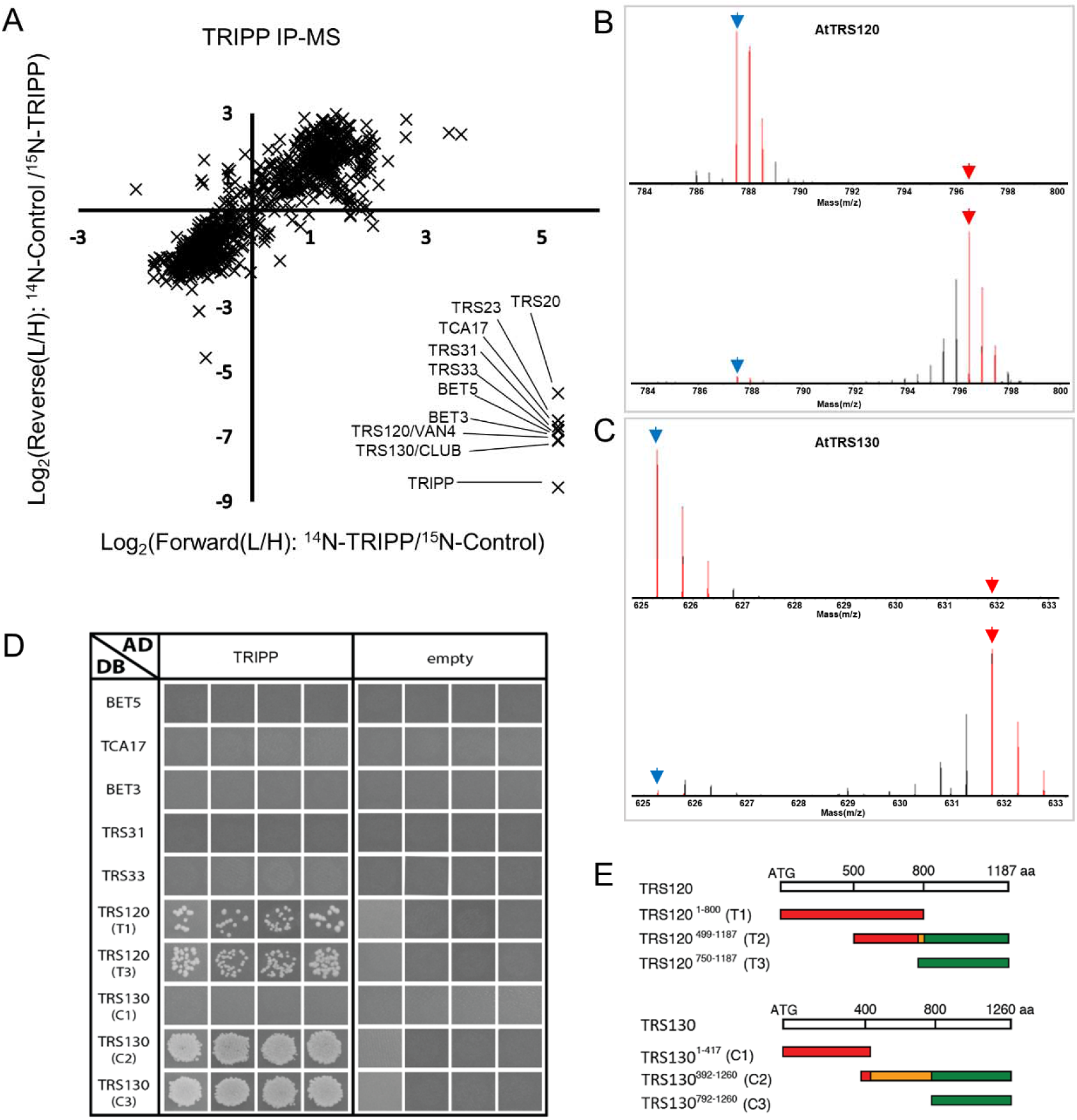
TRIPP interacts with TRAPPII subunits. **A.** Metabolic stable isotope labeling immunoprecipitation mass spectrometry (SILIP-MS) analysis of TRIPP identifies only TRAPPII subunits. SILIP-MS was performed using *35S:TRIPP-YFP*/*tripp-1* complemented plants. Plot shows log_2_ ratio of signal intensities between samples labelled with light (L, ^14^N) and heavy (H, ^15^N) isotopes (L/H ratio) for two biological replicates, in which either the Col-0 control (Forward) or the TRIPP-YFP sample (Reciprocal) was labelled with ^15^N. Note that TRAPP-III subunits are not identified in the data set. **B.** Representative spectra of an AtTRS120 peptide quantified in the SILIP-MS experiments of ^14^N-TRIPP-YFP vs ^15^N-Col-0 (upper panel) and ^14^N-Col-0 vs ^15^N-TRIP-YFP (lower panel). Blue and red arrow points to mono-isotopic peak of the ^14^N and ^15^N labeled peptide respectively. **C.** Representative spectra of an AtTRS130 peptide quantified in the TRIPP-YFP SILIP-MS experiments as descried for panel B. **D.** Yeast two-hybrid assays of interactions between TRIPP and TRAPP subunits. The panels are from different plates. Four independent replicate experiments were performed. The results show interactions of TRIPP with both T1 and T3 regions of AtTRS120 and with the plant specific C2/C3_DB AtTRS130 regions. T2_DB is not included as it is an auto activator, as evidenced by colony growth with the empt AD vector, and this precludes our ability to determine whether AtTRS120_T2 interacts with TRIPP. **E.** TRAPPII coding regions used for yeast two-hybrid interaction assays. Segments colored in red are conserved across kingdoms, while those in green are plant-specific. The orange moiety of the C2 segment is poorly conserved across kingdoms. The T2 middle segment corresponds to sequences found to interact with the exocyst in a yeast two-hybrid screen (Rybak et al., 2014).

### TRIPP directly interacts with the TRAPPII-specific but not shared subunits

If TRIPP associates specifically with the TRAPPII complex, it likely interacts directly with TRAPPII-specific subunits instead of the shared core subunits. To test this, we performed yeast two-hybrid (Y2H) assays of binary TRIPP interactions with known TRAPP subunits (Thellmann et al., 2010). For the TRAPPII-specific subunits, truncations were designed via phylogenetic analysis as described (Figure 3E, Steiner et al., 2016b). In quadruplicate pairwise tests, TRIPP did not interact with shared TRAPP subunits, including AtTRS33, but rather interacted specifically with the two TRAPPII-specific subunits AtTRS120 and AtTRS130 (Figure 3D). Notably, strong interactions were found with the plant-specific or poorly conserved domains of AtTRS130 (Figure 3D, 3E). Positive interactions were also observed with both extremities of AtTRS120 (Figure 3D). In conclusion, the plant-specific TRIPP protein interacts predominantly with plant-specific domains of the large TRAPPII-specific subunits.

### TRIPP localizes to TGN/early endosomal compartments in Arabidopsis roots

Previous studies of TRAPPII specific subunits (e.g AtTRS120 and TRS130) show localization to the *trans*-Golgi network/early endosome (TGN/EE) and to the cell plate (Ravikumar et al., 2018). To examine TRIPP subcellular localization, we generated transgenic plants expressing TRIPP-YFP under the 35S promoter in the *tripp-1* null mutant background (mutant to be described later). Roots of transgenic seedlings expressing TRIPP-YFP were analyzed by confocal imaging. TRIPP-YFP exhibited a punctate localization signal that co-localized with FM4-64 (Figure 4A). TRIPP also exhibited a cell plate localization similar to AtTRS120 and AtTRS130, both TRAPPII-specific subunits (supplemental Figure 2A; Jaber et al., 2010; Rybak et al., 2014). Further, co-localization at the cell plate was confirmed between TRIPP-YFP and TRS120-mCherry in plants expressing both tagged proteins (Figure 4B). TRS33, a subunit shared between different TRAPP complexes, exhibited a similar localization to TRIPP (Supplemental Figure 2B). Co-localization of TRIPP and TRS33 was examined by crossing *35S:TRIPP-mCherry* and *35S:TRS33-YFP* plants. In root cells from plants co-expressing both proteins, all TRIPP-mCherry compartments are positive for TRS33-YFP, whereas about 20% of the TRS33-YFP compartments show no TRIPP-mCherry signal (Figure 4C). This is consistent with TRIPP being a component of only the TRAPPII complex and TRS33 being in both TRAPPII and TRAPPIII complexes. Furthermore, TRIPP-YFP localization is disrupted in the *trs33-1* mutant background consistent with TRIPP being a TRAPP interacting component (Figure 4D). As TRAPPII has been functionally linked to Rab-A GTPases in Arabidopsis (Qi et al., 2011; Qi and Zheng 2011; Naramoto et al., 2014), we performed co-localization experiments with the TGN/EE-localized RAB-A2a (Chow et al., 2008). This revealed that TRIPP co-localized with YFP-RAB-A2a (Figure 4E). To confirm TRIPP localization to TGN is not due to TRAPPII potential interaction with RABs, we perform co-localization with another TGN/EE resident Vesicle transport v-SNARE 12 (VTI12) (Geldner et. al., 2009). Consistently, we observed TRIPP co-localization with VTI12 (Supplemental Figure 2C). Taken together, the data provides *in vivo* evidence that TRIPP co-localizes with TGN/EE-markers and associated TRAPPII subunits (Qi et al., 2011; Naramoto et al., 2014; Ravikumar et al., 2018).

**Figure 4.**
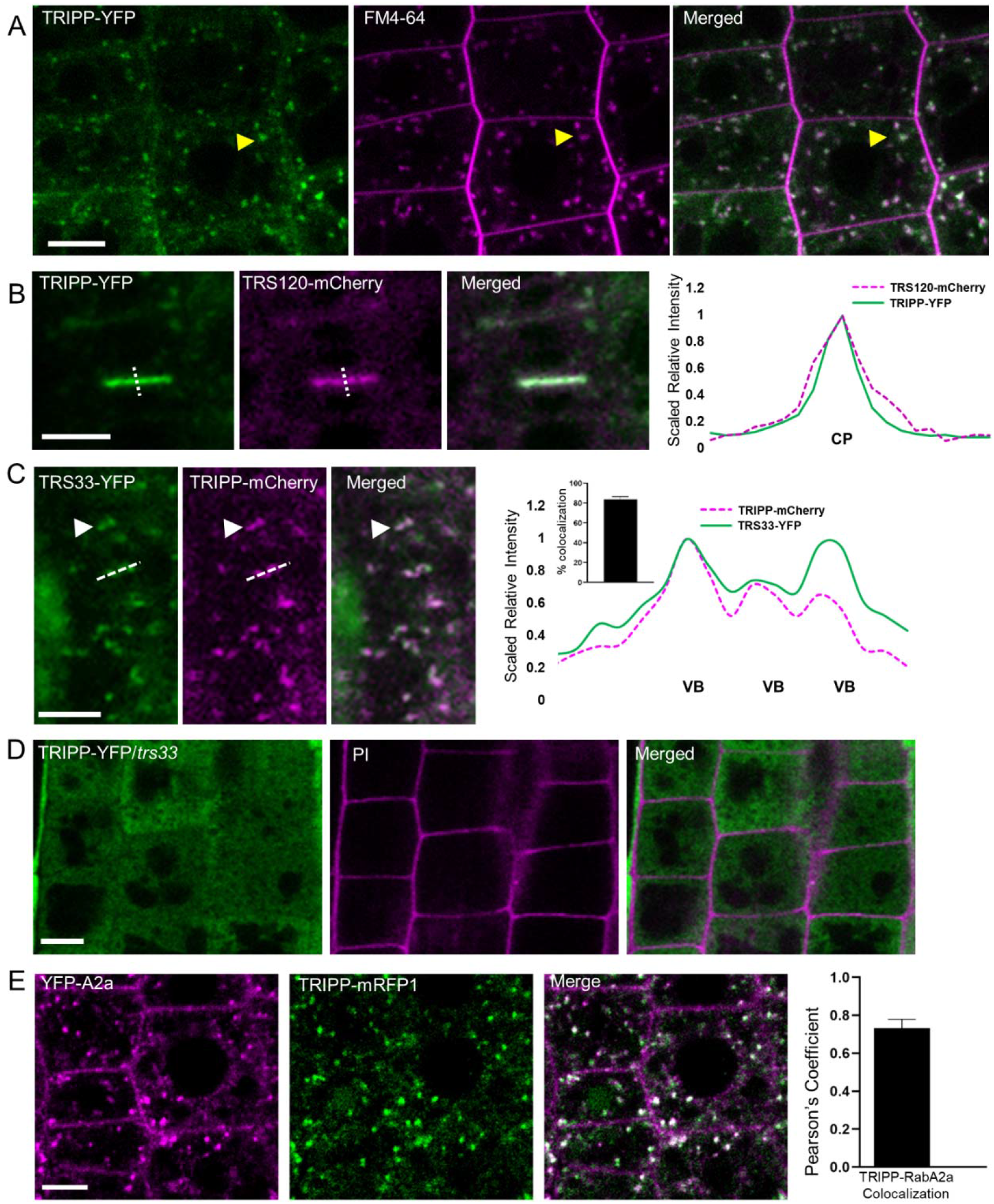
TRIPP localizes to vesicles and the cell plate, similar to known TRAPP components, and its localization disrupted in *trs33-1* mutant background. **A)** Confocal images show TRIPP-YFP and FM4-64 staining signals in the roots of the *35S:TRIPP-YFP*/*tripp* complemented transgenic plants. Arrowhead indicates co-localization of TRIPP-YFP with FM4-64 positive TGN/endocytic vesicles. **B)** TRIPP-YFP co-localizes with TRS120-mCherry to cell plates. Dashed line show the position of the intensity plot. CP = cell plate. **C)** Roots of plants transformed with both *35S:TRS33-YFP* and *35S:TRIPP-mCherry* show co-localization (white arrow head) of TRIPP to some but not all vesicles in which TRS33-YFP localizes to. Dashed line in the image show the position of the quantification graph on the right panel. VB: vesicle bodies, Inset graph shows percent colocalization of TRIPP-mCherry to TRS33-YFP-containing compartments. n = 7. **D)** TRIPP-YFP localization is disrupted in the *trs33-1* mutant background. Root cell walls stained with propidium iodide. **E)** *UBQ10:TRIPP-mRFP1* transgenic plants show TRIPP-mRFP1 colocalization with the TGN marker YFP-RabA2a (Chow et al., 2008). Graph shows co-localization quantification analyzed by Pearson’s correlation coefficient. n = 20 cells. Scale bar for all panels = 5 μm

### The Arabidopsis *tripp* mutant displays growth defects including partial photomorphogenesis in the dark

To explore TRIPP’s role in plant development, we performed phenotypic characterization of *tripp-1*, an insertion allele where a T-DNA is inserted in the 5^th^ exon of the *TRIPP* gene, which has 10 exons and renders a null mutation (Figure 5A, Supplemental Figure 3A, 3B). The *tripp-1* homozygous plants exhibited smaller rosettes compared to wild type (WT) and were sterile (Figure 5B, 5C). Plants heterozygous for the T-DNA insertion did not exhibit a phenotype, and thus the *tripp-1* mutation was maintained in heterozygous plants. To confirm that the phenotypes observed were due to the specific mutation in the *TRIPP* gene, a transgene containing the 35S promoter driving a full-length TRIPP coding sequence fused to YFP was introduced by Agrobacterium transformation to the TRIPP +/− heterozygous plants. Plants expressing TRIPP-YFP in the *tripp-1* homozygous background fully rescued the phenotypes observed in the *tripp-1* mutant (Figure 5), confirming that the observed phenotypes were due to loss of function of TRIPP. Both mutant and WT bolted at similar times but inflorescences were prominently shorter in the mutant (Figure 5C). Floral organs appeared normal in the mutant, with the exception that anthers failed to produce pollen, thus no seeds were produced (Fig 5D). The anthers of the *tripp-1* mutant had a smooth surface suggesting pollen development defects occur in early stages of anther development. To determine whether the *tripp-1* mutant has any defects in the female gametophyte, WT pollen was used to pollinate *tripp-1* mutant flowers. Although the mutant stigma appeared normal, flowers pollinated with WT pollen did not produce any seeds, suggesting that *tripp-1* has defects in both male and female reproductive development. Overall, the *tripp-1* mutant exhibits similar sterility but weaker dwarfism phenotypes compared to the *trs33-1* mutant (Figure 5E-5G). Taken together, our genetic analyses indicate that TRIPP has important functions throughout plant growth and development and particularly for reproductive development.

**Figure 5.**
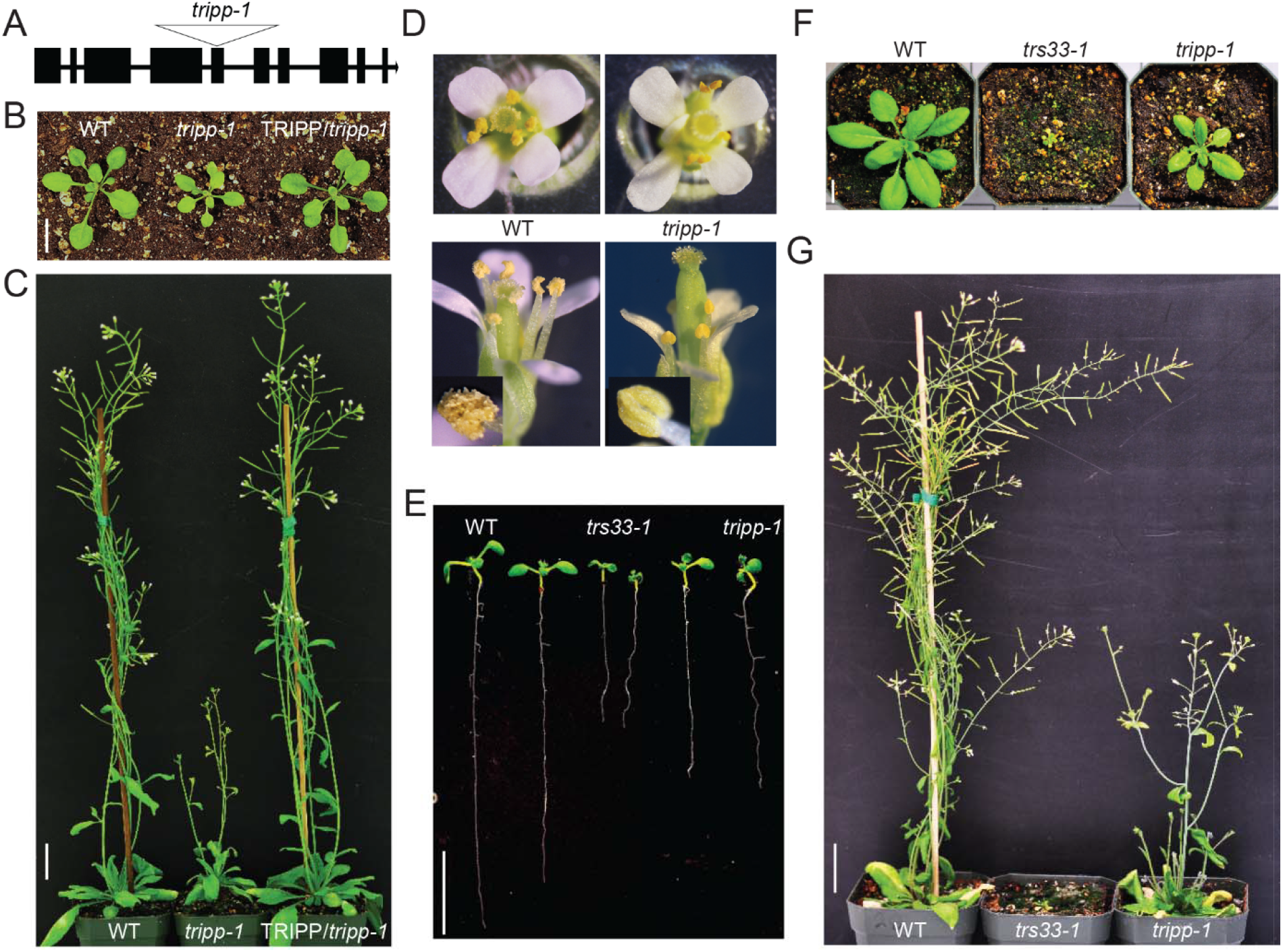
The *tripp* mutant exhibits dwarfism and sterility. **A.** Gene model of Arabidopsis *TRIPP*/AT3G17900 gene shows the T-DNA insertion (Salkseq_063596) of *tripp-1* located in the fifth exon. **B.** Two-week old *tripp-1* mutant plants grown in 16hr light / 8hr dark cycles show smaller rosette compared to WT and the *35S:TRIPP-YFP*/*tripp-1* plants (*TRIPP-YFP*/*tripp-1)*. Scale bar = 1 cm. **C.** Five-week old *tripp-1* mutant plants grown in 16hr light / 8hr dark cycle exhibit significantly shorter inflorescence compare to WT and the *TRIPP-YFP/tripp*-1 plants. Scale bar = 1 inch. **D.** Flowers of *tripp-1* plants appear normal, but anthers lack pollen. Insets of lower panel show magnified images of anthers. **E.** and **F.** The *tripp-1* mutant plants shows shorter roots and stunted growth but phenotype is weaker than *trs33-1* mutant. Scale bar = 1 cm. **G.** Six-week old plants show both *tripp-1* and *trs33-1* mutants exhibit complete sterility phenotypes. Scale bar = 1 inch.

The dwarf phenotype of light-grown *tripp-1* mutant led us to compare *tripp-1* growth and cell elongation in the dark versus light (skotomorphogenesis vs photomorphogenesis). When seeds were placed on medium in the light, seeds germinated at the same time, hypocotyl length was similar between *tripp-1* and WT, but *tripp-1* mutant had shorter roots (Figure 6A-6C). Conversely, in the dark, although the *tripp-1* mutant and WT germinated at the same time, the *tripp-1* mutant exhibited a shorter hypocotyl but longer root compared to WT (Figure 6D-6F). Furthermore, the *tripp-1* mutant grown in the dark for two-weeks developed true leaves, while the WT did not (Supplemental Figure 4). To determine if cell elongation and anisotropy are disrupted in *tripp-1*, the hypocotyl cells of dark-grown and light-grown seedlings were visualized with propidium iodide staining. In the dark, *tripp-1* exhibited prominently shorter cells than WT, whereas light-grown *tripp-1* seedlings showed similar cell size as WT (Figure 6G and 6H). These observations indicate that TRIPP plays an important role in photomorphogenesis.

**Figure 6.**
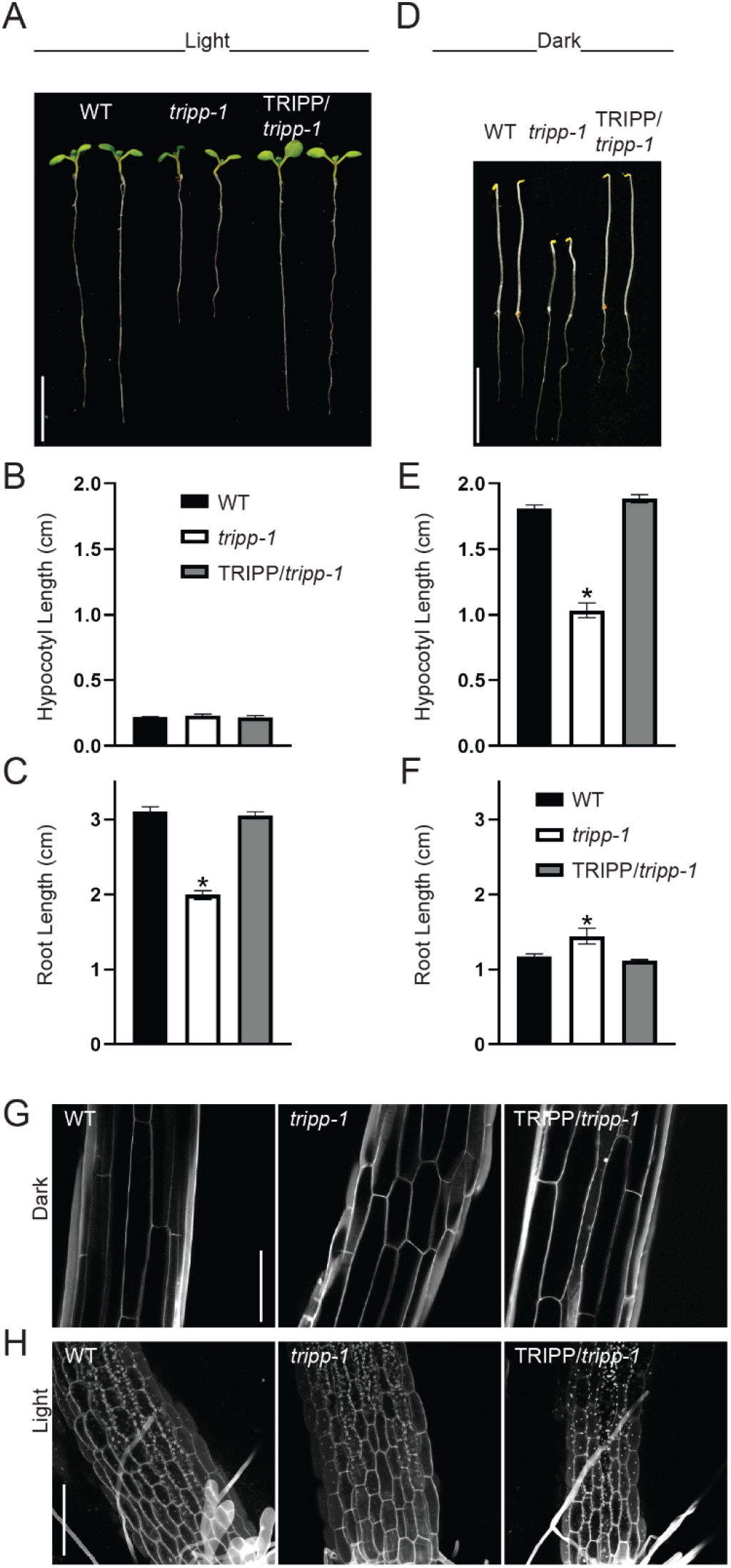
The *tripp-1* mutant exhibits photomorphogenic developmental defects. **A.** Seven-day old *tripp-1* mutant seedlings grown is 16hr light / 8hr dark, show normal hypocotyl length but shorter roots. Scale = 1 cm **B.** and **C.** Quantitation of hypocotyl and root lengths in seven-day light (6hr light/8hr dark) grown seedlings (n=10). Data reported as means ± SEM *P <0.05 compared to WT control (t-test). **D.** Five-day old *tripp-1* mutant seedlings grown in the dark show shorter hypocotyls and longer roots than WT. Scale bar = 1cm **E.** and **F.** Quantitation of hypocotyl and root lengths in 5-day dark-grown seedlings (n=10). Data reported as means ± SEM *P <0.05 compared to WT control (t-test). **G.** Hypocotyl of five-day dark grown *tripp-1* mutant seedlings stained with propidium iodide show a disruption in cell elongation. Scale bar 150 μm. Imaged at hypocotyl elongated zone (lower section of hypocotyl). **H.** Hypocotyl of seven-day light grown seedlings, stained with PI show similar cell growth between *tripp-1* and WT. Scale bar 150 μm. hypocotyl-root junction at bottom section of image.

### TRIPP is present in the green lineage with the exception of chlorophytes

Sequence analysis indicate TRIPP is present in streptophytes (multicellular algae to flowering plants) (Supplemental Figure 5A). TRIPP homologues are found as far as the charophycean algae, which are multicellular photosynthetic organisms ancestral to land plants, but surprisingly not present in unicellular photosynthetic chlorophytes such as chlamydomonas. This is in contrast to other TRAPP components such as the shared TRAPP subunit TRS33, which is present from chlorophytes to embryophytes and across kingdoms (Supplemental Figure 5B; Kim et al., 2016). Protein alignment of TRIPP protein sequence indicates high conservation among its homologues (Supplemental Figure 5C). These analyses indicate that TRIPP might have evolved early in the streptophytes, parallel to the emergence of multicellularity in plants.

## Discussion

The endomembrane system is crucial for cellular organization and morphogenesis, and is regulated by complex networks of proteins and protein-complexes in eukaryotes. The TRAPP complexes have been extensively characterized in yeast and metazoans, and have been shown genetically to play important roles in the endomembrane system in plants, yet the protein components of the TRAPP complex and their functions in plant development remain to be characterized in plants. Our quantitative IP-MS experiments demonstrate that in Arabidopsis TRAPP complexes contain homologues of all thirteen mammalian TRAPP subunits. In addition, the TRAPPII complex associates with an additional plant specific component, which plays important roles in a range of developmental processes including photomorphogenesis.

### The Arabidopsis TRAnsport Protein Particle (TRAPP) complexes include fourteen subunits

Proteomic approaches provide a valuable tool for the characterization of protein-protein interactions and protein complexes in various organisms (Takac et al., 2017). Surprisingly, all previous proteomic studies combined have only identified six TRAPP components in Arabidopsis, in contrast to 11 and 13 subunits in yeast and animals, respectively (Drakakaki et al., 2012; Rybak et al., 2014). For instance, in metazoans, the TRAPPII complex is composed of seven ‘shared’ subunits and three TRAPPII-specific subunits (Figure 6; Scrivens et al., 2011; Riedel et al., 2018; Sacher et al., 2018). While TRAPPIII contains the same shared subunits as TRAPPII, plus four TRAPPIII-specific subunits, including two metazoan-specific not found in yeast (Figure 7; Scrivens et al., 2011; Zhao et al., 2017). Based on our understanding of TRAPP structure in yeast and animals, we focused on using an essential shared subunit combined with quantitative mass spectrometry to identify TRAPP-interacting proteins. TRS33 is a shared TRAPP subunit in yeast and animals, and in Arabidopsis our initial data indicated that AtTRS33 is required for the membrane association of TRS120 (a component of TRAPPII) and for its localization dynamics during cytokinesis (Figure 1). Combined with our earlier findings that *trs33-1* mutants are, like *trappii* mutants, impaired in cytokinesis (Thellmann et al., 2010), this established a function of AtTRS33 as a key component of the TRAPP complexes.

**Figure 7.**
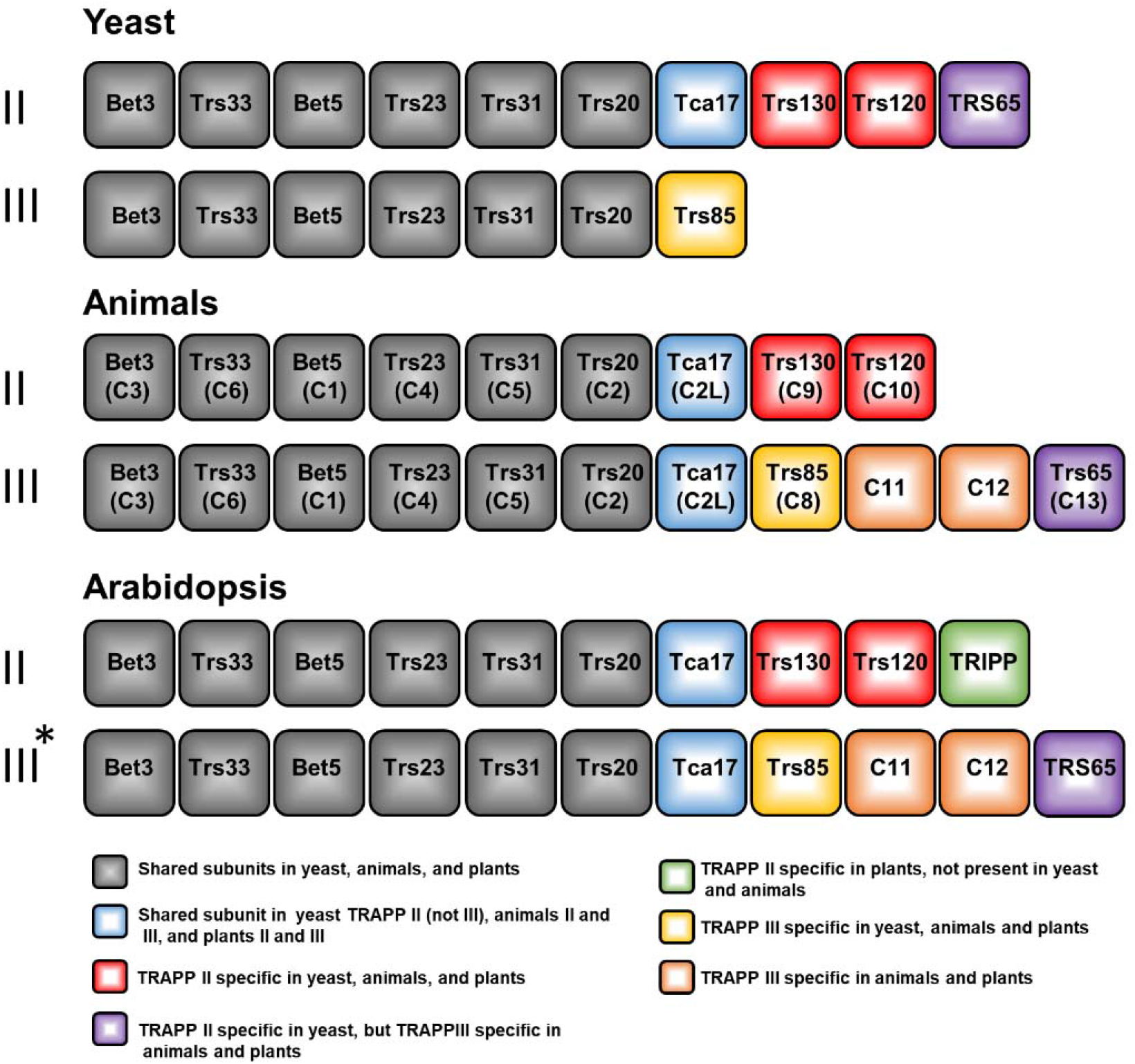
The subunit composition of TRAPP complexes in yeast, animals and Arabidopsis. Overview of the shared and specific subunits for TRAPPII and TRAPPIII present in different kingdoms, with the current report supporting similar TRAPP complex composition in plants. For simplicity original yeast subunit nomenclature is used and in parenthesis the names given to animal homologs for each subunit or only animal-specific names if not present in yeast. Yeast and animal TRAPPII and TRAPPIII complexes have been well defined in previous studies (Thomas et al., 2018; Riedel et al., 2018; reviewed by Kim et al., 2016; Sacher et al., 2018). TRAPP II in plants is defined from the TRIPP IP-MS results presented in this study (Table II). *TRAPPIII in plants is suggested based on the observation that AtTRS33 pulls down all TRAPP subunits (Table I) whereas TRIPP pulls down only the TRAPPII complex but not the TRAPPIII-specific-subunits (Figure 3A and supplemental TABLE II), using the animal orthologous TRAPPIII complex as reference.

In contrast to previous studies (Drakakaki et al., 2012; Rybak et al., 2014), our quantitative IP-MS analysis of AtTRS33 identified fourteen TRAPP interacting proteins, which include homologs of all the TRAPP components known in yeast and animals, as well as a plant-specific protein, TRIPP. We carried out multiple SILIP-MS experiments, each including two replicates, with shoot and root tissues of transgenic plants expressing TRS33 from either the native *TRS33* promoter or the 35S promoter, as well as with different affinity tags. Although more than 1000 proteins were detected by mass spectrometry, accurate quantitation enabled by stable isotope labelling identified only the TRAPP subunit homologs and TRIPP as TRS33-associated proteins. No other proteins showed over 2-fold enrichment by TRS33 in both replicate experiments, and no RAB was found enriched in even a single experiment. It is likely that our stringent washing condition prevented detection of dynamic interactors. But this supports, on the other hand, that TRIPP is a relatively stable component of the TRAPP complex. As such, our high-quality SILIP-MS data identified a more complete set of TRAPP complex components, some of which have not been previously reported to be present in plants. These proteins include homologs of all yeast and metazoan TRAPP subunits, and TRIPP as a new component that has homologs in multicellular photosynthetic organisms but not in fungi or metazoans.

### TRIPP is a plant-specific TRAPPII-associated component

Among studies of TRAPP proteins in different organisms, it is known that animals possess two additional TRAPP subunits (TRAPPC11 and TRAPPC12) compared to yeast (Kim et al., 2016; Riedel et al., 2018). This suggests that multicellular organisms might have evolved a more complex regulation of, or novel functions for, TRAPP components. The human TRAPPC12, for example, has been implicated in kinetochore assembly and mitosis through association with chromosomes (Milev et al., 2015). Interestingly, among the fourteen Arabidopsis TRAPP interactors, TRIPP was found to be specific to plants and is currently classified as an unknown protein in plant databases. Protein sequence analysis and database mining indicate that, based on protein homology, TRIPP is specific to multicellular photosynthetic organisms (Supplemental Figure 6A). This is in contrast to other TRAPP subunits such as TRS33, which is present in all eukaryotes including unicellular algae and flowering plants (Supplemental Figure 5B).

Our biochemical and genetic data strongly support that TRIPP is specifically associated with TRAPPII, a complex that has been shown to be involved in post-Golgi membrane trafficking (Ravikumar et al., 2018). While the SILIP-MS analysis of AtTRS33 interactome identified fourteen subunits, the TRIPP SILIP-MS identified only ten subunits that include the specific subunits of TRAPPII but exclude those of TRAPPIII. The presence of the homologs of the four animal TRAPPIII-specific subunits (TRAPPC8, TRAPPC11, TRAPPC12, and TRAPPC13) in the AtTRS33 interactome but not the TRIPP interactome suggests the existence of both TRIPP-containing TRAPPII complex and an equivalent of TRAPPIII complex that excludes TRIPP. Indeed, in accordance with our results, a recent IP-MS using AtTRAPPC11 has confirmed the presence of a TRAPPIII complex in Arabidopsis (Rosquette et al., 2019). Our Y2H data indicates that TRIPP interacts with the two TRAPPII-specific subunits TRS120 and TRS130, further corroborating TRIPP association with the TRAPPII complex. Consistent with these molecular interactions, TRIPP co-localizes with AtTRS33 at some endosomal compartments presumably in the TRAPPII complex, but is absent at a subset of AtTRS33-containing endosomal compartments, which are likely TRAPPIII-specific (Figure 4C). These results support that TRIPP is a specific component of TRAPPII.

The TRAPPII complex is well conserved in yeast and animals, with the exception of the TRS65 subunit, which in yeast associates with TRAPPII and in animals with TRAPPIII (Choi et al., 2011). Now we show that TRAPPII is also conserved in plants but, as in animal models, TRS65 does not associate with TRAPPII. Thus, plant TRAPP complexes seem more similar to metazoan than to yeast complexes. Our results demonstrate that multi-cellular plants have evolved TRIPP as a plant-specific component of the TRAPPII complex, whereas TRAPPIII is conserved in animals and plants (Figure 7).

TRAPPII is known to associate with the TGN, which is a hub for membrane trafficking; exocytosis, endocytosis and protein sorting (Qi et al., 2011; Naramoto et al., 2014; Ravikumar et al., 2018). Arabidopsis TRAPPII subunits, TRS120 and TRS130, localize to endosomes, the *trans*-Golgi network, and cell plates during cytokinesis (Ravikumar et al., 2017; Ravikumar et al., 2018). Our study shows that TRIPP exhibits a similar localization to TGN/EE compartments as well as cell plate. The disruption of TRIPP localization in the *trs33* mutant supports that TRIPP localization to membrane compartments is dependent on the TRAPP complexes.

### TRIPP plays critical roles in plant growth, reproduction and light-dependent development

The *tripp-1* mutant exhibits a range of developmental phenotypes, including partial dwarfism, shorter roots, and sterility in light-grown plants. The sterility phenotype is similar to that observed in the *trs33-1* mutant (Supplemental Figure 4). Mutations of the TRAPPII specific components in plants cause detrimental phenotypes (Thellmann et al., 2010; Rybak et al., 2014; Ravikumar et al., 2017). The overall growth phenotypes of *tripp-1* are weaker than the *trs33*, or *trs120* and *trs130* mutants, which have lethal phenotypes past the early seedling stage. These observations suggest that TRIPP association with TRAPPII provides specific functions, in contrast to the broader functions of AtTRS120 and AtTRS130. As TRIPP is only present in multicellular photosynthetic organisms, while other TRAPPII components are present in all eukaryotes including algae, it is conceivable that TRIPP may play important roles in development and/or environmental adaptation in multicellular plants. Indeed, the phenotypes of *tripp* mutants grown in the dark suggest a role in photomorphogenesis (Figure 6). While membrane trafficking is critical for proper cell growth, and any defect in vesicle traffic regulation can cause hypocotyl growth defects (He et al., 2018), the dark-grown *tripp-1* mutant seedlings exhibit not only shorter hypocotyls but also longer roots, a hallmark of photomorphogenesis. These results demonstrate that TRIPP, and likely the TRAPPII complex, plays a role in light-regulated plant development. Our proteomic and genetic study provides a framework for future studies of TRAPP complex functions in cellular membrane trafficking and plant development.

## METHODS

### Plant materials

Arabidopsis thaliana ecotype Columbia (Col-0) was used in this study. Plants were grown in greenhouses with a 16-h light/ 8-h dark cycle at 22°C for general growth and seed harvesting. For seedling analysis such as dark germination or for root imaging, plants were grown on petri dishes as follows: seeds were surface-sterilized with 70% ethanol+0.1%Triton X-100 for 5 mins, dried on a filter paper and then grown on ½ Murashige and Skoog (MS) medium, 1% sucrose and 0.8% phytoagar. Plates with seeds were stratified at 4 °C for 3 days and then placed in a growth chamber under 16hr light/8hr dark (or complete dark for germination in the dark) at 22 °C.

T-DNA insertional mutant, *tripp-1* (SALKseq_063596.3) was obtained from the Arabidopsis Biological Resource Center (www.arabidopsis.org; Alonso et al., 2003). *trs33-1* (SALK_109244) is from our previously published work (Thellmann et al., 2010).

### Construction of plasmid and generation of transgenic plants

Full-length coding sequences for AtTRS33 and TRIPP without stop codons were cloned into *pENTR/SD/D-TOPO* (Invitrogen). To generate *35S-TRS33-YFP* and *35S-TRIPP-YFP* constructs, each entry clone was sub-cloned into Gateway-compatible destination vector *pEarleyGate-101* using LR clonase (Invitrogen). To generate *35S-TRIPP-mCherry*, entry clone was subcloned into *pEarleyGate 101* vector that carries the mCherry tag instead of YFP. For native promoter driven TRS33-4XMyc-6XHis, an entry clone was generated using a 1kb genomic region upstream of TRS33 start site fused with TRS33 full coding sequence and sub-cloned into a gateway compatible *pCAMBIA1390* vector. For co-localization with YFP-RabA2a, TRIPP was cloned from Arabidopsis (Columbia) genomic DNA into *pUB-RFP-DEST-HygR* (derived from Grefen et al., 2010). Constructs were validated by nucleotide sequencing. All binary vector constructs were introduced into Agrobacterium tumefaciens (strain GV3101) and transgenic plants generated Arabidopsis by floral dipping (Clough and Bent, 1998). *TRS120:TRS120-GFP* and *PUBQ:TRS120-mCherry* are described in Rybak et al., 2014 and *YFP-Rab-A2a* in Chow et al., 2008.

### ^15^N Stable-isotope-labeling in Arabidopsis quantitative MS analysis of the AtTRS33 interactome

The pTRS33:TRS33-MycHis and wild-type plants were grown for two weeks at 22°C with 24 hours light on vertical ^14^N or ^15^N medium plates (½ Murashige & Skoog modified basal salt mixture without nitrogen 0.39g/L, 8g/L phytoblend, and 1g/L KNO_3_ or 1g/L K^15^NO_3_ (Cambridge Isotope Laboratories) for ^14^N medium or ^15^N medium, respectively, pH5.8). About 2.5 g of tissue was harvested for each sample, ^14^N-labeled TRS33-Myc-His and ^15^N-labeled wild type samples or reciprocal (^14^N-labeled wild type and ^15^N-labeled TRS33-Myc-His) were mixed, then ground in liquid nitrogen and stored in −80°C. Immunoprecipitation was performed as followed; proteins were extracted in 10 mL NEBT buffer (20 mM HEPES, pH7.5, 40 mM KCl, 1 mM EDTA, 10% glycerol, 0.5% Triton X-100, 1 mM phenylmethylsulfonyl fluoride (PMSF), Pierce protease inhibitor cocktail (Thermo Fisher), and PhosStop cocktail (Roche)), centrifuged at 5,000rpm for 5 min, supernatant (SP) transferred to new tube and centrifuged a second time for 15 min at 14,000 rpm, then SP transferred to new tube. The SP was incubated with 10μl of Myc-Trap®_MA (chromotek) and incubated for 1 hour. Next, beads were washed 3X with 1ml immunoprecipitation buffer without detergent. Proteins were eluted with 2X SDS buffer. The eluted proteins were separated by SDS-PAGE. After Coomassie Brillant blue staining, the protein bands were excited and subjected to in-gel digestion with trypsin.

The peptide mixtures were desalted using C18 ZipTips (Millipore) and analyzed on a Q-Exactive HF hybrid quadrupole-Orbitrap mass spectrometer (Thermo Fisher) equipped with an Easy LC 1200 UPLC liquid chromatography system (Thermo Fisher). Peptides were separated using analytical column ES803 (Thermo Fisher). The flow rate was 300nL/min and a 120-min gradient was used. Peptides were eluted by a gradient from 3 to 28% solvent B (80% acetonitrile/0.1 formic acid) over 100 mins and from 28 to 44% solvent B over 20 mins, followed by short wash at 90% solvent B. Precursor scan was from mass-to-charge ratio (m/z) 375 to 1600 and top 20 most intense multiply charged precursor were selection for fragmentation. Peptides were fragmented with higher-energy collision dissociation (HCD) with normalized collision energy (NCE) 27. Tandem mass spectrometry peak lists were extracted using an in-house script PAVA, and data were searched using Protein Prospector (Chalkley et al., 2008) against the *Arabidopsis* Information Resource (TAIR10) database, to which reverse sequence versions were concatenated (a total of 35,386 entries) to allow estimation of a false discovery rate (FDR). Carbamidomethylcysteine was searched as a fixed modification and oxidation of methionine, Peptide N-terminal Gln conversion to pyroglutamate and N-terminal acetylation as variable modifications. Data were searched with a 10 ppm tolerance for precursor ion and 20 ppm for fragment ions. Peptide and protein FDRs were set as 0.01 and 0.05. ^15^N labeled amino acids were also searched as a fixed modification for ^15^N data. Quantification was done using Protein Prospector.

Note that for the IP-MS of p35S-TRS33-YFP vs wild-type or p35S-TRIPP-YFP vs wild-type the same method described above was used with the difference that GFP-trap magnetic agarose beads (Allele) were used instead for immunoprecipitation.

### Yeast Two-Hybrid (Y2H)

Y2H screens were performed as described (Dreze et al., 2010). TRAPPII-specific (AtTRS120 and CLUB/AtTRS130) truncations are described in Steiner et al., 2016; shared TRAPP subunits were obtained from Riken full length cDNA clones(Seki et al., 2002) or from a collection of 12,000 Arabidopsis ORFs(Wessling et al., 2014). We used the GAL4 DNA-binding domain (DB) encoded in Y2H vector pDEST-pPC97, subsequently transformed into the yeast strain Y8930, as well as gene fusions to the Gal4 activation domain (AD) in the yeast strain Y8800 (Altmann et al., 2018). Both strains are derived from PJ68-4 (James et al., 1996). The constructs were screened by yeast mating in quadruplicate pair-wise tests. Screening was done as a binary mini-pool screen, i.e. each DB-ORF was screened against pools of 188 AD-ORFs. Interactions were assayed by growth on selective plates using the HIS3 reporter, and using 1mM 3-Amino-1,2,4-triazole (3-AT) to suppress background growth. This primary screen was carried out once and interaction candidates were identified by Sanger sequencing. All candidate interactions were verified by pairwise one-on-one mating in three independent experiments. Only pairs scoring positives in all three assays were considered as bona fide interaction partners.

### Confocal microscopy

Plant roots were imaged on an Olympus (www.olympus-ims.com) Fluoview 1000 confocal laser scanning microscope (CSLM), a Leica TCS SP8 microscope (www.leica-microsystems.com) and a Zeiss 880 CLSM (www.zeiss.com/microscopy). Imaging data were acquired using LAS-X software (Leica) and professed on Fiji/image J software (www.imagej.com). The following excitation and emission parameters were used for the respective fluorescent proteins: YFP 514nm excitation, 525-575nm emission; GFP 488nm excitation, 500-550nm emission; mCherry 561nm excitation, 580-643nm emission; mRFP 561nm excitation, 581-754nm emission; and FM4-64 emission at 700-725 nm. For Figure 1, cell cycle stages were determined via TRS120-GFP localization dynamics, taking into account how this membrane marker follows phragmoplast microtubule dynamics (Steiner et al., 2016a). As *trapp* mutants affect both cell plate biogenesis and phragmoplast microtubule dynamics (Steiner et al., 2016a), we also relied on FM4-64 to image cell plates in *trapp* mutants. Based on the appearance of the cell plate, cytokinesis was broadly split into early stages (cell plate initiation and biogenesis) versus late stages (cell plate insertion and maturation; (Smertenko et al., 2017). For quantification of co-localization analysis, image background was subtracted from both channels; in the resultant background-subtracted images, co-localization between intracellular YFP: RAB-A2a and TRIPP: mRFP1 was quantified using Pearson’s Correlation Coefficient using the JACoP Fiji plugin (Costes et al., 2004; Bolte and Cordelieres, 2006).

### Accession numbers

Sequence data from this article can be found in the GenBanK/EMBL data libraries under the following accession numbers: TRS33, AT3G05000; TRIPP, AT3G17900; BET5, AT1G51160; TRS20, AT1G80500; TCA17, AT2G20930; BET3, AT5G54750; TRS23, AT5G02280; TRS31, AT5G58030; TRS85, AT5G16280; TRS120, AT5G11040; TRS130, AT5G54440;TRAPPC11, AT5G65950;TRAPPC12, AT4G39820;TRAPPC13, AT2G47960; RAB-A2a, At1g09630.

All data that support the conclusions of this study are available from the corresponding author upon request.

## Supplemental Information

Supplemental Table 1. 35S-TRS33-YFP SILIP-MS results from roots and shoots.

Supplemental Table 2. 35S-TRIPP-YFP SILIP-MS results with additional information such as peptide number and fold change.

Figure S1. Mass Spectrum of a TRIPP peptide identified in TRS33-MycHis SILIP-MS.

Figure S2. TRIPP cell plate localization and TRS33 localization and additional TGN marker colocalization.

Figure S3. *tripp-1* T-DNA genotyping and mRNA analysis by RT-PCR

Figure S4. Two-week dark grown *tripp-1.*

Figure S5. TRIPP phylogenetic and protein alignment.

## AUTHOR CONTRIBUTIONS

V.J.G. performed all experiments, with the exception of those mentioned below, and wrote the manuscript.

Z.W. conceived the research project, designed experiments, and edited the manuscript.

S.X. and W.W. performed mass spectrometry analyses.

R.R. generated, analyzed and assembled data shown in Figure 1 under the supervision of F.F.A.

L.E., M.F., and I.M. provided constructs and performed the experiment shown in Fig. 4E.

L.E. edited the manuscript.

M.A. and R.R. performed yeast-two-hybrid experiments in Figure 3 under the supervision of F.F.A and P. F-B.

F.F.A. designed experiments, performed crosses, analyzed data, drafted sections of and edited the manuscript.

Z.W., F.F.A., S.X., and P.F-B. acquired funding.

## Acknowledgments

We are very thankful to Alma Burlingame and Robert Chalkley for assistance with mass spectrometry data analysis using Protein Prospector, Heather Cartwright from the Carnegie Imaging Facility for technical assistance with microscopy, Adam Idoine for help with phylogenetic analysis and revising manuscript, Miriam Abele for critically appraising the literature and for supporting us with assembling figure panels, as well as Andreas Czempiel for technical assistance. We thank the WZW/TUM Centre for Advanced Light Microscopy (CALM) for access to confocal microscopes, and Roman Meier at the TUMmesa facility, directed by Sharon Zytynska, for supporting us with optimal growth conditions for our plants.

