## Supplemental Data for "Proteomic studies of the Arabidopsis TRAPP complexes reveal conserved organization and a novel plant-specific component with a role in plant development"

Garcia et al. Plant Cell SUPPLEMENTAL FIGURES

| TRAPP subunits identified by 35S-TRS33-YFP immunoprecipitation from Arabidopsis roots | | | | | | | |
| --- | --- | --- | --- | --- | --- | --- | --- |
|  | **Forward** | | | **Reciprocal** | | |  |
| **Gene ID** | **Peptide No** | **% Cov** | **L/H ratio** | **Peptide No** | **% Cov** | **L/H ratio** | **Protein Name** |
| AT1G51160 | 7 | 44.4 | 2.63 | 3 | 18.9 | 0.40 | BET5 |
| AT1G80500 | 4 | 43.7 | 2.98 | 3 | 28.9 | 0.25 | TRS20 |
| AT2G20930 | 13 | 88.6 | 4.11 | 5 | 57.1 | 0.24 | TCA17 |
| AT5G54750 | 33 | 66.1 | 27.90 | 8 | 43.5 | 0.02 | BET3 |
| AT5G02280 | 8 | 59.6 | 2.77 | 3 | 26.2 | 0.30 | TRS23 |
| AT5G58030 | 6 | 34.9 | 3.45 | 2 | 7.7 | 0.18 | TRS31 |
| AT3G05000 | 24 | 87.9 | 16.60 | 3 | 19.1 | 0.05 | TRS33 |
| AT5G16280 | - | - | - | - | - | - | TRS85 |
| AT5G11040 | 19 | 18.5 | 2.79 | 8 | 7.8 | 0.36 | TRS120 |
| AT5G54440 | 23 | 19.3 | 4.56 | 7 | 6.8 | 0.26 | TRS130 |
| AT5G65950 | 25 | 26.2 | 11.90 | 22 | 23.5 | 0.08 | TRAPPC11 |
| AT4G39820 | 7 | 14.5 | 4.43 | 8 | 22.1 | 0.20 | TRAPPC12 |
| AT2G47960 | 11 | 23.5 | 2.72 | 6 | 14.0 | 0.34 | TRS65 |
| AT3G17900 | 23 | 29.1 | 2.66 | 9 | 10.6 | 0.36 | TRIPP |
| TRAPP subunits identified by 35S-TRS33-YFP immunoprecipitation from Arabidopsis Shoots | | | | | | | |
|  | **Forward** | | | **Reciprocal** | | |  |
| **Gene ID** | **Peptide No** | **% Cov** | **L/H ratio** | **Peptide No** | **% Cov** | **L/H ratio** | **Protein Name** |
| AT1G51160 | 9 | 74.0 | 4.52 | 2 | 14.2 | 0.45 | BET5 |
| AT1G80500 | 4 | 37.8 | 3.63 | 1 | 8.9 | 0.35 | TRS20 |
| AT2G20930 | 16 | 90.7 | 6.31 | 7 | 64.3 | 0.25 | TCA17 |
| AT5G54750 | 38 | 66.1 | 53.50 | 6 | 38.7 | 0.01 | BET3 |
| AT5G02280 | 6 | 53.9 | 4.42 | 2 | 14.2 | 0.34 | TRS23 |
| AT5G58030 | 9 | 42.1 | 7.87 | 2 | 10.3 | 0.31 | TRS31 |
| AT3G05000 | 40 | 90.2 | 25.00 | 2 | 14.5 | 0.02 | TRS33 |
| AT5G16280 | 2 | 1.5 | 25.90 | - | - | - | TRS85 |
| AT5G11040 | 23 | 20.7 | 4.36 | 4 | 3.3 | 0.38 | TRS120 |
| AT5G54440 | 60 | 43.9 | 9.13 | 2 | 1.7 | 0.21 | TRS130 |
| AT5G65950 | 35 | 34.2 | 14.50 | 16 | 16.1 | 0.11 | TRAPPC11 |
| AT4G39820 | 5 | 14.5 | 8.72 | 2 | 4.9 | 0.23 | TRAPPC12 |
| AT2G47960 | 16 | 35.5 | 4.72 | 2 | 4.5 | 0.15 | TRS65 |
| AT3G17900 | 35 | 45.3 | 3.68 | 11 | 13.6 | 0.43 | TRIPP |

**Supplemental Table I. Fourteen Arabidopsis TRAPP components were identified by stable isotope labeling immunoprecipitation mass spectrometry (SILIP-MS) using *35S:TRS33-YFP*.** SILIP-MS was performed using the transgenic plants expressing TRS33-YFP under the 35S promoter (Root and Shoot separately). Number of unique peptides (Peptide No.), percentage of sequence coverage (% Cov), and ratio of signal intensities between samples labelled with light (L, ^14^N) and heavy (H, ^15^N) isotopes (L/H ratio) are shown for two biological replicates, in which either the Col-0 control (Forward) or the TRS33-YFP sample (Reciprocal) was labelled with ^15^N.

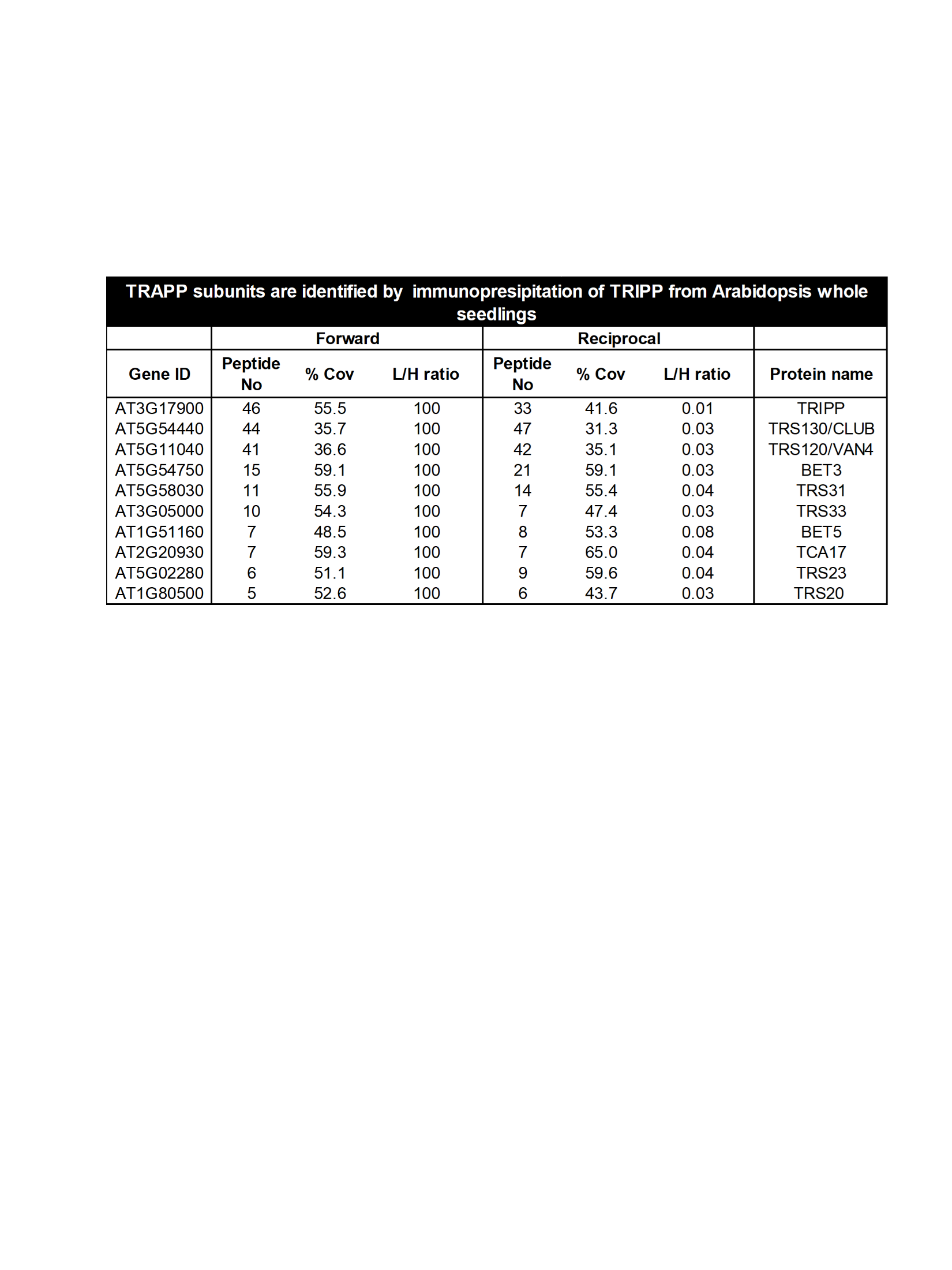

**Supplemental Table II. Metabolic stable isotope labeling immunoprecipitation mass spectrometry (SILIP-MS) analysis of TRIPP identifies only TRAPPII subunits.** SILIP-MS was performed using *35S:TRIPP-YFP*/*tripp* complemented plants**.** Numbers of unique peptides (Peptide No.), percentage of sequence coverage (% Cov), and ratio of signal intensities between samples labelled with light (L, ^14^N) and heavy (H, ^15^N) isotopes (L/H ratio) are shown for two biological replicates, in which either the Col-0 control (Forward) or the TRIPP-YFP sample (Reciprocal) was labelled with ^15^N.

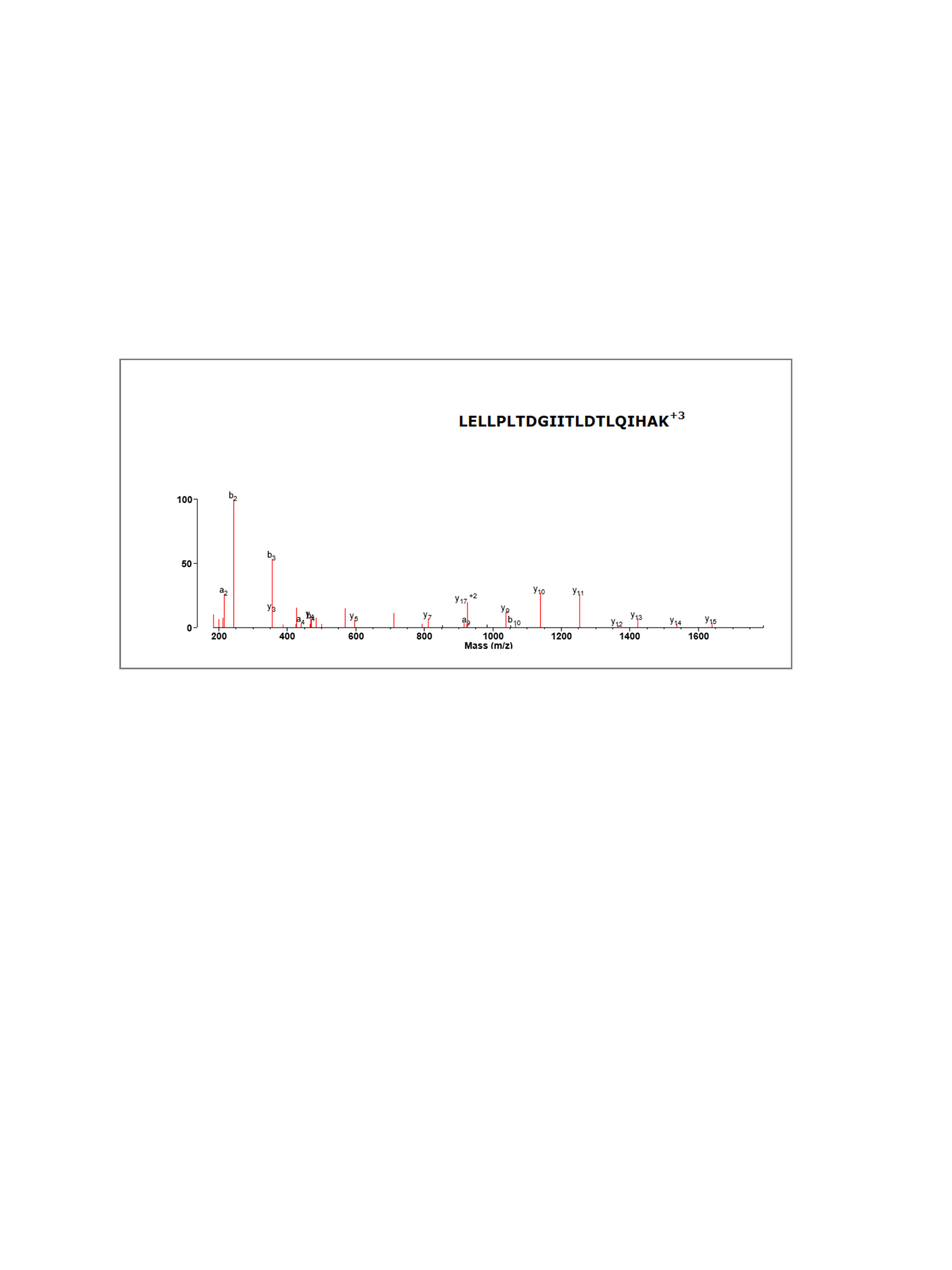

**Supplemental Figure 1.** HCD mass spectrum of a peptide from TRIPP spanning from amino acid 793-813.

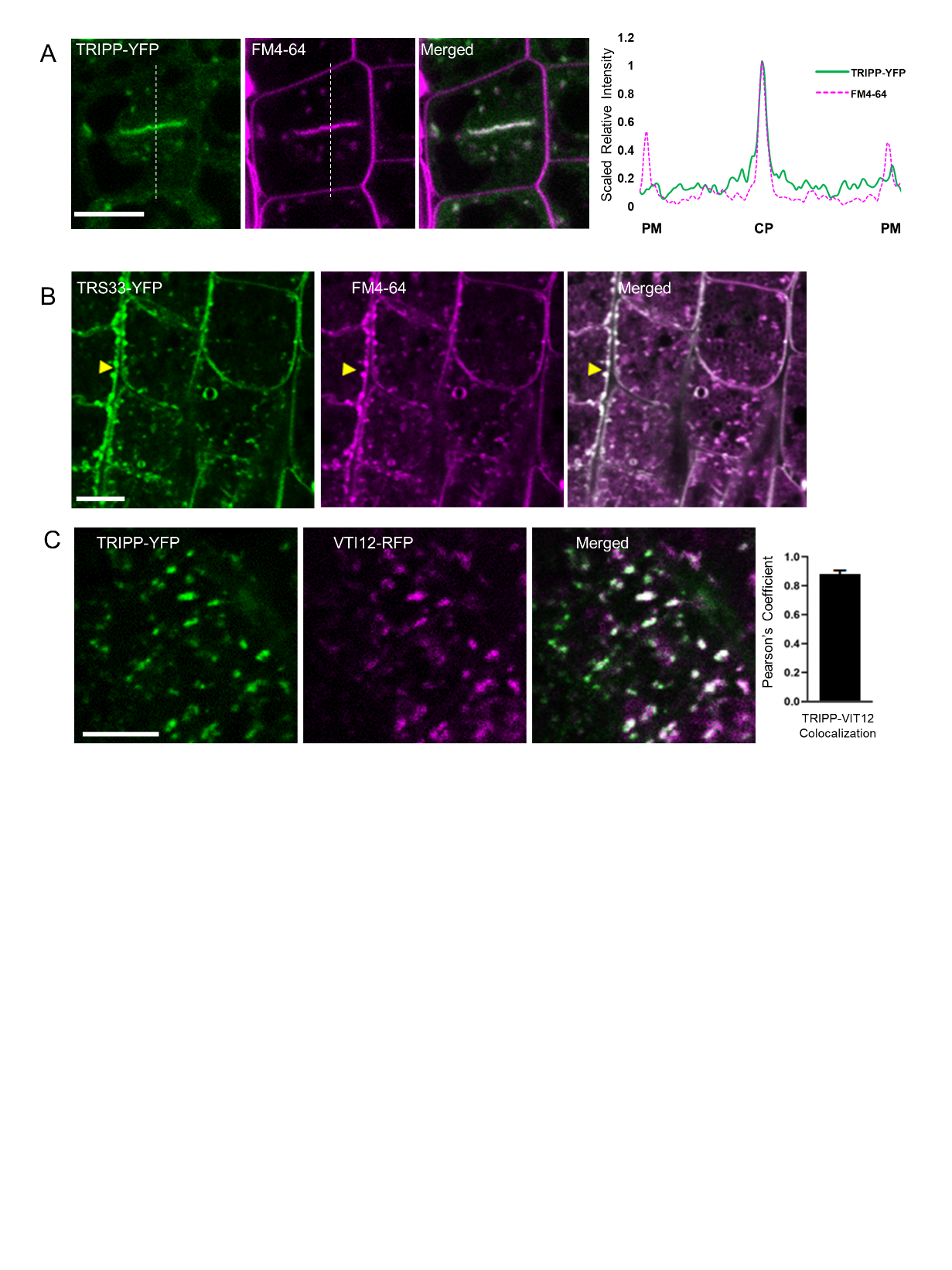

**Supplemental Figure 2. TRIPP co-localizes with TRAPP components**

**A.** TRIPP-YFP localizes to cell plate. Dashed lines how the position of the intensity plot. CP = cell plate, PM = plasma membrane.

**B.** TRS33-YFP exhibits punctate and co-localization with FM4-64, in the *35S:TRS33-YFP* plants. Scale bar = 5µm.

**C.** TRIPP-YFP co-localizes with TGN resident VTI12. Scale bar = 5µm.

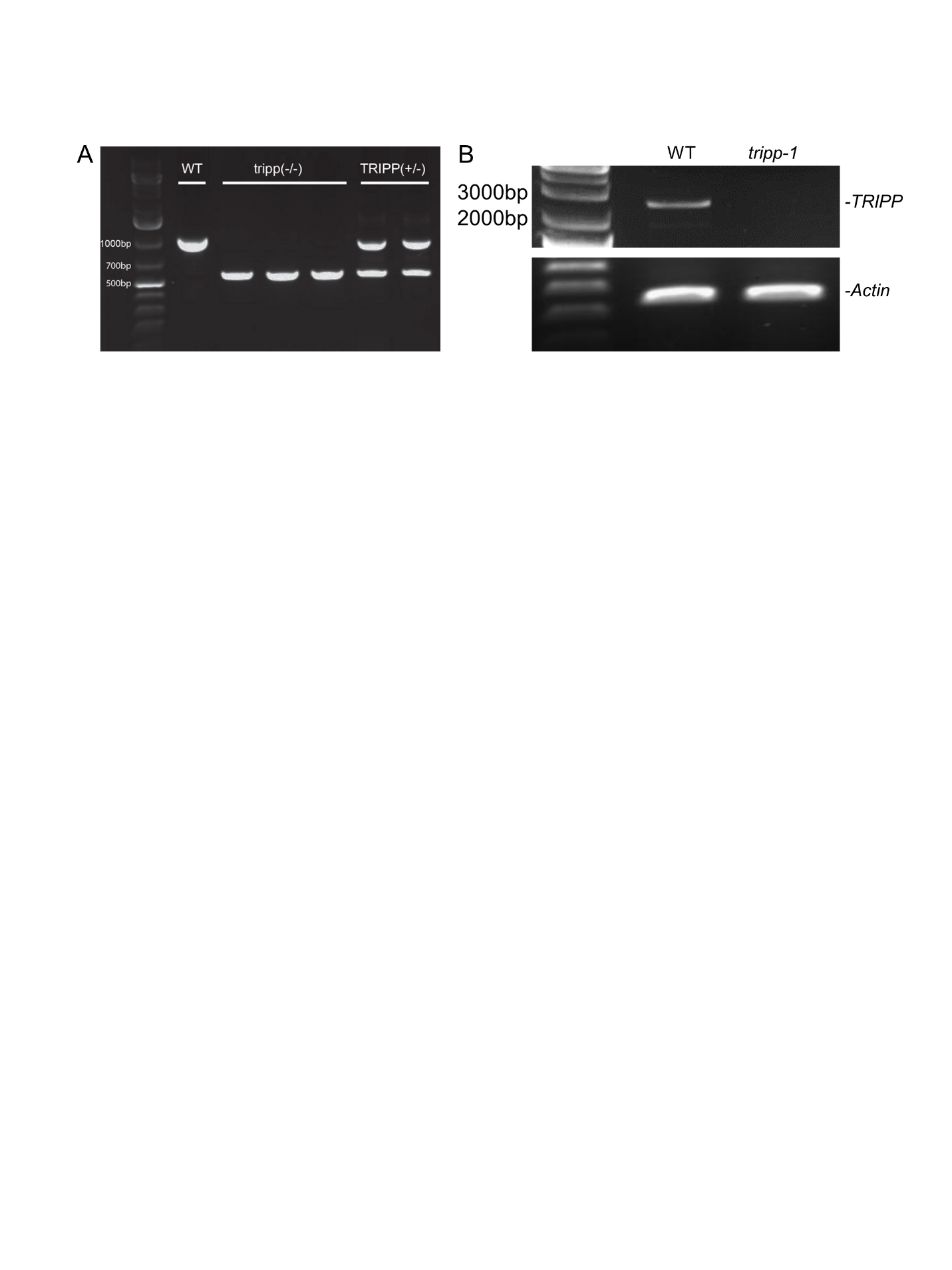

**Supplemental figure 3. Genotyping of *tripp-1* (SALKseq_063596.3 T-DNA insertion).**

**A.**Six representative plants genotyped with LP+RP+LBb1.3 primers combined. WT plant shows only upper band indicating WT DNA amplification, T-DNA homozygous plants show only lower band indicating amplification of T-DNA fragment, whereas heterozygous plants can amplify both WT and T-DNA fragments.

**B.** RT-PCR analysis indicate *tripp-1* is KO.

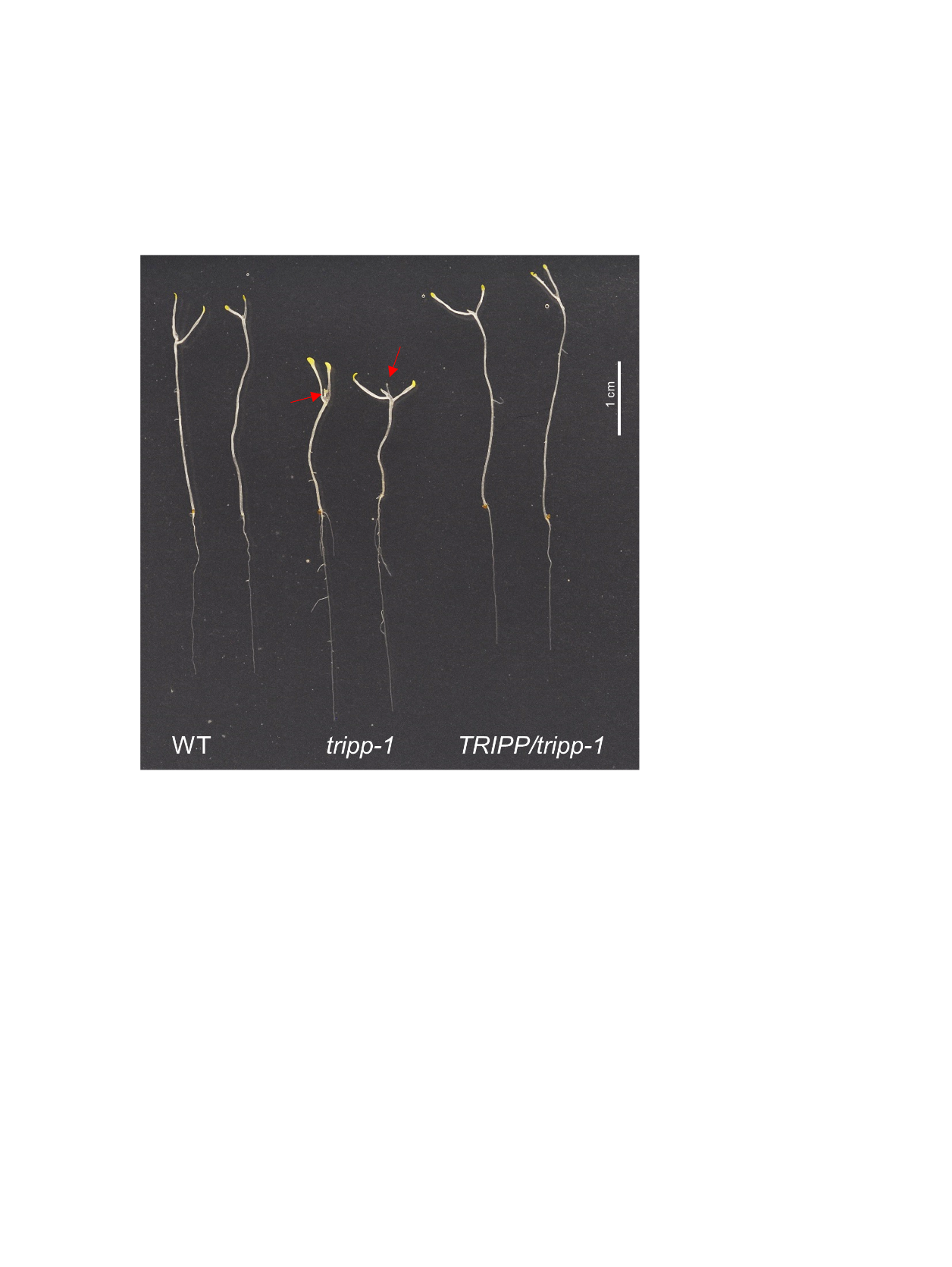

**Supplemental figure 4. Two-week dark grown *tripp-1* mutant exhibits light grown developmental phenotype.** *tripp-1* develops true leaves in long-term dark conditions (red arrows).

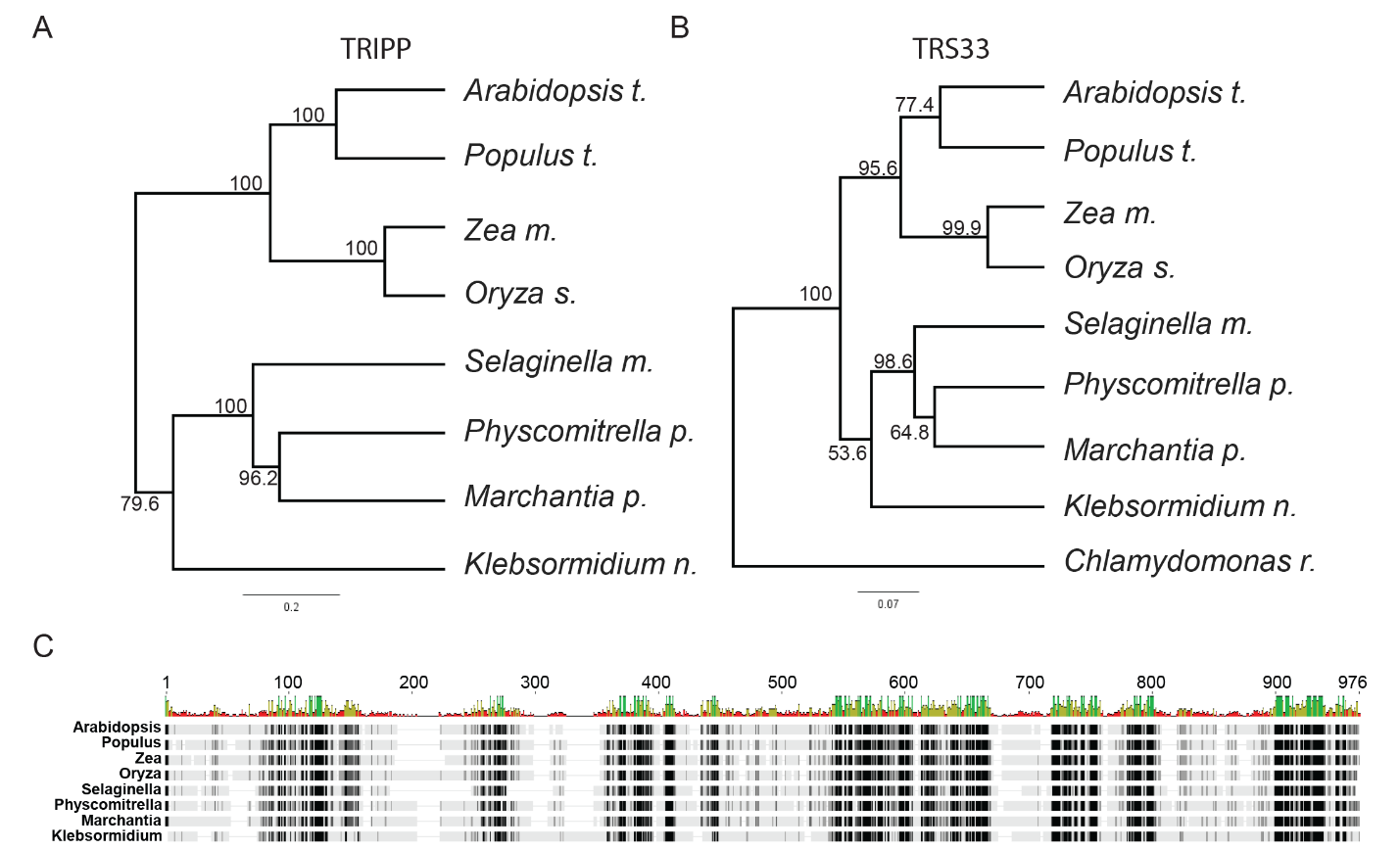

**Supplemental figure 5. TRIPP is present in multicellular photosynthetic organisms**

**A**. Phylogenetic tree representing TRIPP in the green lineage, with homologs found only in streptophytes (multicellular algae to flowering plants) but absent in chlorophytes such as the unicellular algae chlamydomonas. Selected representative species are shown in tree.

**B**. Phylogenetic tree representing TRS33, a shared TRAPP subunit, present in all organisms in the green lineage (as well as across kingdom, not shown in tree) in contrast to TRIPP.

**C**. TRIPP alignment with representative homologues from different plant taxa showing high conservation. Black residues show 100% similarity, dark grey 80-100% similarity, light grey 60-80% similarity, and white represents residues with less than 60% similarity.

**Supplemental Methods:**

**Genotyping and RT-PCR analysis of *tripp-1***

Genomic DNA was extracted from T-DNA insertion mutant *tripp-1*, WT, and complemented plants as followed. Leaf tissues were homogenized in genomic DNA extraction buffer 200mM Tris-Cl, pH 7.5, 250 mM NaCl, 25 mM EDTA, and 0.5% SDS and used to identify a region flanking the T-DNA insertion site by PCR. Primers used TRIPP-1 LB, TRIPP-1 RB, and T-DNA primer LBb1.3. RNA extraction was performed with Sigma-Aldrich Spectrum Plant Total RNA Kit. *TRIPP* cDNA was synthesized using revertAid reverse transcriptase enzyme from Thermo Fisher Scientific according to instructions.

**Protein alignment and Phylogenetic tree**

Proteins homologous to TRIPP were aligned using the ClustalW algorithm in the Geneious software suite. Alignments were manually adjusted for optimization. No protein with significant homology to AtTRIPP was found in chlorophytes. Positions are highlighted according to their percent similarity based on the BLOSUM62 matrix. The Accessions used were, *Arabidopsis thaliana*: NP_566591.1, *Chlamydomonas reinhardtii*: not found, *Zea mays*: XP_020403649.1, *Physcomitrella patens*: XP_024399605.1, *Populus trichocarpa*: XP_024466417.1, *Oryza sativa*: XP_015629884.1, *Selaginella moellendorffii*: XP_024528913.1, *Marchantia polymorpha subsp. ruderalis*: OAE18401, *Klebsormidium nitens*: KFL_000080290. To generate the TRIPP phylogenic tree, we use distance tree of selected homologous TRIPP protein sequences, generated using Geneious Tree Builder, using the Jukes-Cantor genetic distance model and UPGMA tree building method. Bootstrapping 100000X. Accessions were as for the TRIPP alignment. Whereas for generation of TRS33 phylogenetic tree, the same methods as TRIPP were used with the following accessions: *Arabidopsis thaliana*: NP_187151, *Chlamydomonas reinhardtii*: XP_001701459.1, *Zea mays*: NP_001149190.1, *Physcomitrella patens*: XP_024372395, *Populus trichocarpa*: XP_024456893, *Oryza sativa*: XP_015612457.1, *Selaginella moellendorffii*: XP_002966358, *Marchantia polymorpha subsp. ruderalis*: PTQ49625, *Klebsormidium nitens*: GAQ89824.
